# Characterizing the landscape of gene process dependencies in cancer

**DOI:** 10.1101/2025.11.14.688518

**Authors:** Julia B. Curd, Navin Kumar Balagopal, Adam L. Green, Gregory P. Way

## Abstract

Precision oncology aims to tailor cancer treatment to tumor genetics but it currently benefits only a small fraction of patients, in part because the primary focus is to match drug targets to single genes. The Cancer Dependency Map Project (DepMap) aimed to characterize the landscape of single-gene dependencies, which increased the universe of potential drug targets. However, the common challenges of drug off-target effects and polypharmacology may limit effectiveness of single genes as drug targets. To address this limitation, we apply BioBombe, an AI/ML framework, to DepMap gene dependency data. This approach characterizes gene process dependencies, which are groups of genes within a biological process that cells rely on for survival. BioBombe fits many hundreds of dimensionality reduction models, across a large range of latent dimensionalities. We find that this multiple-model approach discovers many more gene process dependencies than any single model alone. Using Reactome and CORUM-based gene set enrichment analyses, we characterize the landscape of gene process dependencies, identifying, for example, mitotic regulation and the citric acid cycle as targets, as well as many cancer type-specific dependencies. In gliomas, for example, TP53-and mitochondrial-related pathways emerged as key process vulnerabilities. Linking gene process dependencies with drug sensitivity scores on matched cell lines, we discovered both established and novel drug candidates. Furthermore, we developed a machine learning approach to predict gene process dependencies and associated drug sensitivities from RNA-seq data. Using this approach, we predicted cladribine as a potential new therapeutic for pediatric high-grade glioma and validated its effectiveness at killing these cells. Taken together, BioBombe provides a scalable and interpretable framework for uncovering complex gene process dependencies, which guides drug repurposing, and introduces a novel targeting paradigm for precision oncology.

## Introduction

Precision oncology, which tailors’ patient treatment based on the unique genetic profiles of individual tumors [1], has emerged as a powerful approach for improving outcomes for cancer patients. While this strategy has produced notable successes, such as the development of targeted inhibitors against EGFR mutations in lung cancer [2] and BCR-ABL fusions in chronic myeloid leukemia [3], it remains limited in scope. Fewer than 10% of cancer patients currently benefit from genetically-guided treatment strategies [4], and only about 15% of human proteins are considered druggable [5]. Large-scale initiatives such as The Cancer Dependency Map Project (DepMap) [6] have accelerated the discovery of single-gene vulnerabilities, leading to clinical trials for targets like PRMT5 [7]. Yet focusing narrowly on individual genes overlooks the reality that cancer is much more complex. Cancer is driven by coordinated, multi-gene processes spanning signaling pathways, protein complexes, and metabolic networks [8]. A deeper understanding of these higher-order dependencies may translate precision oncology into broadly effective therapies.

The potential of multi-gene targeting is also underscored by clinical successes. PARP inhibitors, for example, exploit synthetic lethal interactions within DNA damage repair pathways [9], while immune checkpoint blockade leverages microsatellite instability to amplify anti-tumor immunity [10]. These cases highlight the therapeutic power of targeting coordinated biological processes rather than isolated genes. Recent advances in CRISPR-Cas9 screening approaches further demonstrate the utility of genome-scale perturbation data for identifying survival-essential genes, drug resistance mechanisms, and therapeutic vulnerabilities relevant to precision oncology [11]. Still, such successes remain exceptions, and the overall landscape of multi-gene dependencies is poorly defined. The DepMap Achilles CRISPR knockout project provides genome-scale perturbation data across hundreds of cancer cell lines, designed to uncover gene dependencies and synthetic lethalities [6]. However, related analyses treat these data at the level of individual genes, leaving hidden process-level signals underexplored. A framework that can systematically extract and interpret biological processes from these gene dependency screens would address this gap.

Recent work has begun to address this gap by moving beyond single-gene dependencies to model the functional architecture underlying large-scale gene dependency datasets. For example, Webster introduced a framework for modeling pleiotropy in genetic fitness screens, recognizing that a single gene perturbation may influence multiple biological functions or pathways [12]. By representing each gene’s effect as a combination of independent, empirically-derived functional components, Webster disentangles overlapping signaling pathways, identifies distinct roles for multifunctional genes, and predicts complex stoichiometries from fitness data. Furthermore, the FIREWORKS framework advanced the analysis of genetic coessentiality, or the tendency of genes with shared biological functions to exhibit correlated fitness profiles across cell lines [13]. FIREWORKS introduced a user-accessible platform for constructing coessentiality networks centered on user-specified genes. Together, Webster and FIREWORKS demonstrated that higher-order functional structures ranging from pleiotropic interactions to coessential gene modules can be computationally recovered from large-scale gene dependency data.

We hypothesize that unsupervised compression algorithms will identify higher-order gene processes, such as pathways, signaling networks, and protein complexes, and will reveal previously unrecognized therapeutic vulnerabilities in cancer. To test this, we extended our original BioBombe approach of training multiple distinct machine learning models across many latent dimensionalities to DepMap gene dependency data (version 24Q2) [14]. Initially applied to RNAseq gene expression data, BioBombe optimizes gene signatures of cell type and gene mutations, and demonstrated that training multiple compression algorithms outperformed any single model. Similar consensus-based and multimodal machine learning approaches have shown improved robustness and predictive performance across biomedical applications, including network-based drug repurposing and community detection analyses, where individual algorithms often produced inconsistent or complementary results [15, 16]. The original BioBombe algorithms included principal component analysis (PCA) [17], independent component analysis (ICA) [18], non-negative matrix factorization (NMF) [19], denoising autoencoders (DAE) [20], and variational autoencoders (VAE) [21]. BioBombe fit these models across several different latent dimensionalities ranging from 2 to 200, and, for the autoencoders, across five different random initializations. Here, we extended the algorithms to incorporate β-variational autoencoders (β-VAEs) and β-total correlation VAEs (β-TCVAEs), which are more recent VAE variant enhancements. β-VAEs introduce a tunable parameter to balance reconstruction fidelity and feature disentanglement [22], while β-TCVAEs penalize correlations between latent variables to promote feature representation independence [23]. We also fit these models across 2 to 200 latent dimensionalities. By analyzing the DepMap gene dependency data through this expanded suite of models, we sought to move beyond single gene-level dependencies to systematically uncover interpretable latent dimensions corresponding to gene process dependencies in cancer. Importantly, this study differs from the original BioBombe application not only in data modality (gene dependency versus transcriptomic profiles) but also in cohort composition and cancer-type representation, factors that may contribute to differences in model performance and biological signal recovery.

Our analyses revealed thousands of significant gene process dependencies across cancer types, including well-established dependencies such as TP53 signaling and mitotic checkpoint pathways, as well as unexpected processes that may represent novel therapeutic opportunities. Importantly, by integrating with PRISM drug screening data [24], we linked these dependencies to drug sensitivity across hundreds of clinically-relevant compounds. This approach enabled both discovery of new higher-order targets and putative repurposing of existing FDA-approved drugs. For instance, we confirmed strong associations between TP53-related gene process dependencies and sensitivity to MDM2 inhibitors [6]. We also developed a machine learning approach to predict gene process dependencies from RNA-seq data and observed variable results: some dependencies manifested strongly in RNA-seq signals while others could not be recovered. We applied this RNA-seq prediction approach to a separate dataset of pediatric high-grade gliomas and molecularly tested predictions for three drugs. All three drugs showed positive correlations with cell killing, but only cladribine was significantly associated with predicted activity in our dataset. Broadly, BioBombe provides a scalable, interpretable framework for discovering and mapping gene process dependencies in cancer. By extending precision oncology from single-gene targets to multi-gene processes, our study establishes a rich foundation for both novel therapeutic discovery and rational drug repurposing, with implications for improving the reach and effectiveness of precision therapies.

## Results

### Applying BioBombe to DepMap gene dependencies

The BioBombe [14] approach fits many different dimensionality reduction models, by training different algorithms and systematically toggling the internal latent dimensionality size *k*. The BioBombe philosophy is to train many different algorithms to find diverse latent, or hidden, representations that represent biologically-meaningful signatures useful for various tasks such as cancer subtyping, characterizing gene expression landscapes of gene mutations, cell typing from bulk gene expression data, and more. Rather than selecting a single model, BioBombe trains many thousands of models, which optimizes these signatures.

Here, we extended the original BioBombe framework, adding more modern VAE variants: β-VAE, and β-TCVAE. Both of these architectures identify so-called disentangled representations, including loss functions that encourage independent latent dimensions. We also dropped the DAE, as it found less optimal signatures in every case compared to the VAE. We trained these six algorithms across twenty-eight latent dimensionalities, ranging from 2 to 200. Another core difference between the original BioBombe and our current implementation, is that rather than training these models on bulk RNAseq gene expression data, we apply BioBombe to DepMap gene dependency data (version 24Q2) (**Fig. 1A**). The 24Q2 DepMap gene dependency data was collected using genome-wide CRISPR knockouts to measure the gene effect on cell viability across 1,150 cell lines and 73 cancer types (**Supplementary Figure 1**). After filtering samples lacking age and cancer type metadata, we applied BioBombe to 933 cell lines (see **Methods** for more details). We hypothesize that applying BioBombe to DepMap gene dependencies will reveal higher-order dependencies representing core biological processes, protein complexes, or other multi-gene patterns. We call these patterns gene process dependencies, which only become apparent when studying relationships across multiple single-gene dependencies.

**Figure 1.**
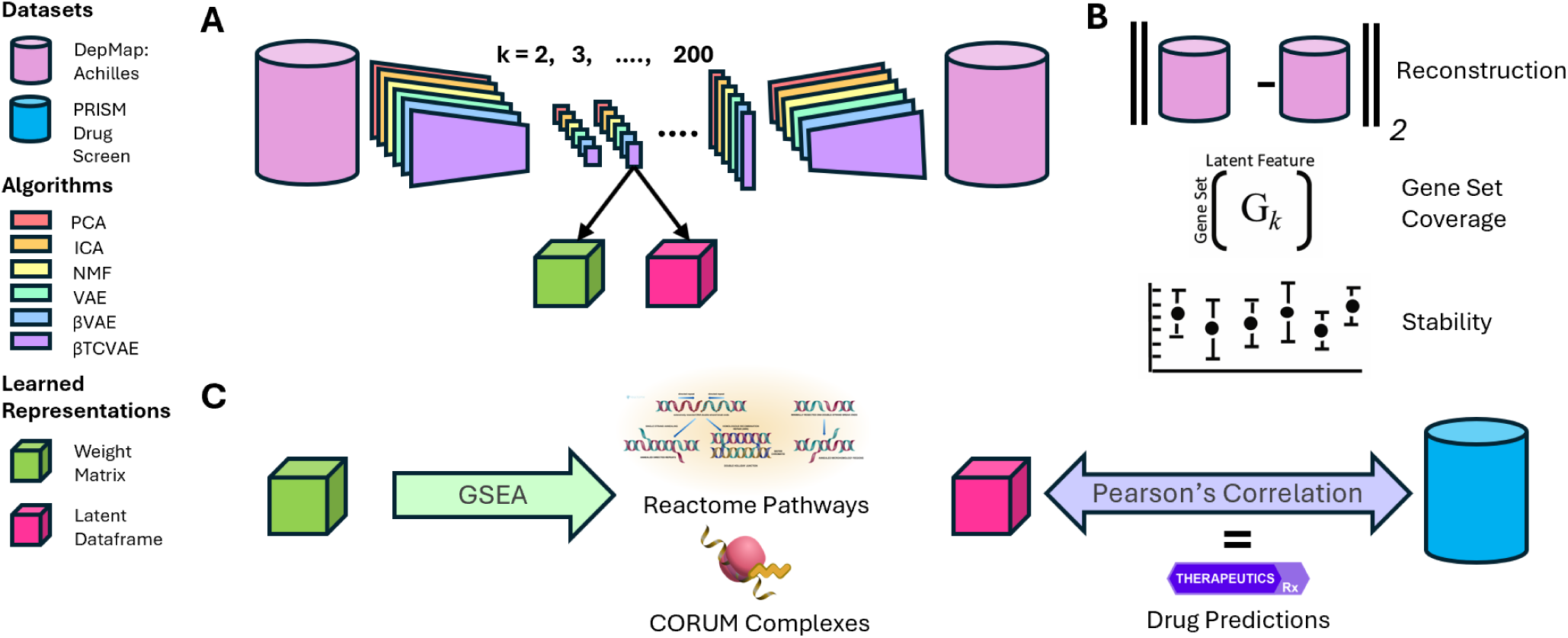
Overview of the BioBombe framework to find, evaluate, and leverage gene process dependencies. **(A)** We apply six machine learning algorithms, including PCA, ICA, NMF, VAE, β-VAE, and β-TCVAE, across a range of latent dimensionalities from *k* = 2-200 to gene dependency data from the Cancer Dependency Map (DepMap) CRISPR knockout screens. Each model fit generates two outputs: a weight matrix (green cube) and a latent representation (z matrix; pink cube). **(B)** We evaluate each model using three different metrics: Reconstruction, gene set coverage, and stability. **(C)** We perform Gene Set Enrichment Analysis (GSEA) on the weight matrix to identify enriched biological processes and interpret the gene process dependencies (Reactome pathways and CORUM protein complexes). Considering matched cell lines, we apply Pearson’s correlation to the latent dataframe with drug sensitivity from the PRISM dataset to identify putative drug matches to specific gene process dependencies.

To train our BioBombe models, we stratified the gene dependency data, balanced by age and sex, into 70 percent training data, 15 percent testing and 15 percent validation. This approach was motivated by the known influence of developmental context and sex-associated biology on gene expression variability across cancer types. Pediatric and adult tumors frequently exhibit distinct molecular programs, while sex-linked transcriptional differences may contribute additional structure captured by latent representations. To evaluate BioBombe performance, we assessed reconstruction accuracy, representation stability, and biological breadth per model fit **(Fig. 1B)**. Reconstruction error, measured using mean squared error (MSE), quantified the difference between input and reconstructed gene dependency profiles. We used Centered Kernel Alignment (CKA) to assess the stability and similarity of models across different initializations and algorithms, and we computed gene set coverage to measure the diversity of biological pathways and protein complexes recovered across models.

After training BioBombe, we extract two key matrices from each model: the weight matrix and the latent dataframe (z matrix). The weight matrix includes the latent dimensions which represent gene process dependencies, and the latent dataframe represent sample-specific scores for these specific gene process dependencies. We analyze these BioBombe results in two key ways. First, for each model, we applied Gene Set Enrichment Analysis (GSEA) [25] to the weight matrix using the Reactome [26] and CORUM (the comprehensive resource of mammalian protein complexes) [27] libraries. We chose Reactome because the library includes thousands of recorded pathways, reactions, and complexes.

Additionally, it performed the best in initial prototyping across library testing with high enrichment scores and many significant results. We selected CORUM due to its extensive verification processes and protein complex information, as well as important cancer relevancy. This GSEA approach adds an interpretive layer to the gene process dependencies we find through BioBombe, interpreting specific pathway and protein complex targets. Second, we correlated small molecule drug sensitivity (derived from matched cell lines based on the PRISM assay [24]) with the latent dataframe for each model (**Fig. 1C**). We use these correlations to develop drug repurposing hypotheses about which small molecules are effective at killing cancer cell lines expressing the specific gene process dependency.

### Prototyping and evaluating BioBombe with stability, reconstruction, and gene set coverage

As we’re applying BioBombe to DepMap gene dependency data for the first time, it is necessary to confirm feasibility. Among the six methods we implemented (PCA, ICA, NMF, VAE, β-VAE, and β-TCVAE), PCA and ICA are deterministic approaches that do not require iterative training, while NMF and the neural network-based models (VAE, β-VAE, and β-TCVAE) are trained to optimize reconstruction objectives. The original BioBombe algorithms (PCA, ICA, NMF, and VAE) were tested extensively, to ensure they could be applied to a different data type than the original model used. High enrichment scores from the VAE models motivated us to explore additional autoencoder architectures in this extension, with the goal of capturing additional biologically significant pathways. Because β-VAE and β-TCVAE are new additions to the BioBombe suite, we focused specifically on demonstrating feasibility with these algorithms. We used the python optimization tool Optuna [28] to select optimal hyperparameters (beta, learning rate, batch size, number of epochs, and optimizer type), initially for a *k* = 50 model and then for models spanning all other latent dimensionalities. Training the prototype *k* = 50 models with optimized hyperparameters, we observed high performance with steadily decreasing loss over time (**Fig. 2A**). The β-VAE displays smoother convergence, with a more stable loss curve, while the β-TCVAE exhibits noisier behavior, with visible fluctuations particularly in later epochs, which may reflect sensitivity to the total correlation regularization or early signs of overfitting. Despite this difference, both models successfully compress the high-dimensional single gene dependency profiles into lower-dimensional latent dimensions. Confident in BioBombe’s ability to model these data, we next optimized the full BioBombe suite, across all latent dimensionalities, applied to DepMap gene dependency data using the same Optuna approach.

**Figure 2.**
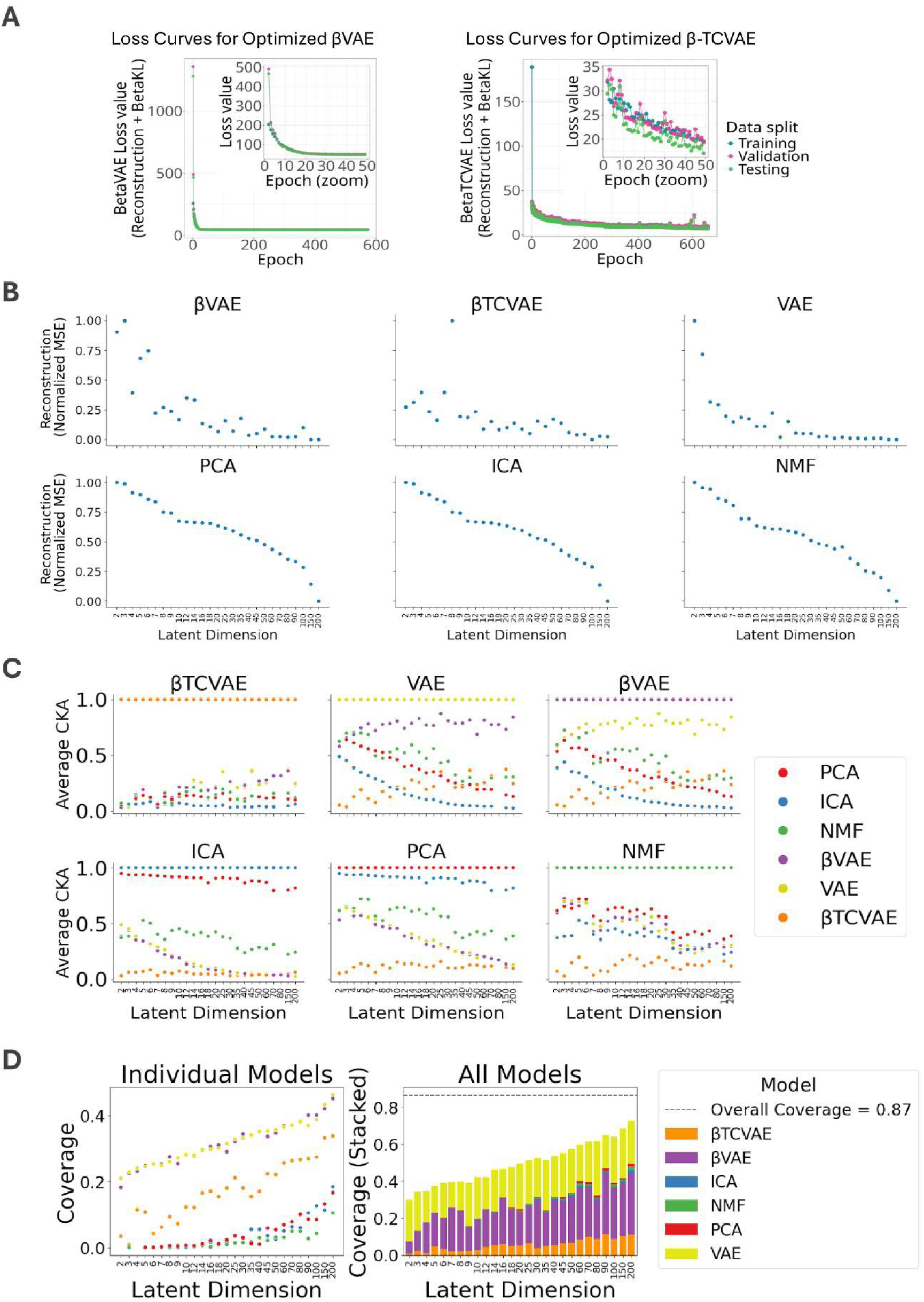
Evaluating BioBombe representations. **(A)** Training, validation, and testing loss curves for βVAE and β-TCVAE models, the two added to the original BioBombe suite, for training a prototype *k* = 50 model on DepMap gene dependency data. Training the full BioBombe suite we report **(B)** Reconstruction Mean Squared Error (MSE) values for each model across the latent dimensionalities. **(C)** Centered Kernel Alignment (CKA) stability comparison between each model across the latent dimensionalities. We observe high similarity between ICA and PCA as well as βVAE and VAE, with large differences across the dimensionalities for other model comparisons. **(D)** Gene Set Coverage, the proportion of total unique pathways or complexes recovered, for each individual model/latent dimension combination and coverage for each dimension separated by model stacked for all models.

To evaluate model performance, we measured both mean squared error (MSE) of the reconstructed output and CKA [29] for stability. The MSE metric quantifies how well models reconstruct the input data, and CKA captures the stability within models and similarity of latent dimensions across different models (across algorithms, latent dimensionalities, and model initializations). We also calculated Gene Set Coverage, which is defined as the proportion of the total number of unique pathways or protein complexes recovered as significantly enriched by Gene Set Enrichment Analysis (GSEA). We assessed reconstruction error for each model, as measured as normalized MSE between input and reconstructed gene dependency profiles (**Fig. 2B**). We calculated (and show) reconstruction scores based on the test dataset. Normalized MSE was consistently lowest in the VAE and β-VAE models, and decreased sharply as latent dimensionalities increased. In contrast, the PCA, ICA, and NMF models showed higher reconstruction errors at all latent dimensionalities, with a similar notable decrease in MSE as the latent dimensionality increased. This is likely due to their inability to capture non-linear gene dependency interactions, which were overcome only at higher latent dimensionalities. Notably, β-TCVAE displayed slightly higher reconstruction error than VAE and β-VAE but still outperformed the linear methods. These findings suggest that VAE architectures are generally better suited for reconstructing complex gene dependency patterns.

Our CKA analysis, also performed on the test dataset, revealed several interesting trends in comparing the latent representation signals of different models and across dimensionalities (**Fig. 2C**). Specifically, when comparing latent dimensions between models, ICA and PCA show the highest CKA similarity, consistent with their shared linear structure. β-VAE and VAE also demonstrate high CKA similarity, reflecting their architectural similarity. In contrast, β-TCVAE shows low CKA similarity with all other models, suggesting that its total correlation regularization induces uniquely structured latent dimensions (**Supplementary Figure 2**). Furthermore, ICA-NMF and PCA-NMF comparisons yield only moderate CKA values with a slightly decreasing trend across latent dimensionalities, but with noticeable jaggedness, suggesting only partial redundancy in the features they capture. In contrast, all other model pairs (e.g., ICA-VAE, PCA-β-VAE, NMF-β-TCVAE) show a strong and consistent downward trend in CKA as latent dimensionality increases. This suggests that as models are allowed more latent capacity, their learned representations diverge, capturing increasingly distinct biological signals.

Within-model CKA values across latent dimensions are generally non-linear for all three VAE models, reflecting this sensitivity to initialization at different latent dimensionalities. In contrast, cross-model CKA comparisons show a smoother, more linear decrease as latent dimensionality increases, highlighting the benefit BioBombe derives from optimizing diverse signatures across both dimensionalities and model architectures.

We also conducted an analysis of gene set coverage. Using a Reactome gene set library, we applied GSEA to all latent dimensionalities (see next section for more details), and assigned latent dimensions to the most significant gene set after correcting for false discoveries. We calculated gene set coverage as the proportion of significant gene sets over the total number of gene sets in the respective library.

When considering Reactome, the coverage results suggest that models with more complex architectures (e.g., VAE, β-VAE and β-TCVAE) tend to uncover a broader range of significant gene sets, potentially due to their capacity to learn more disentangled and non-linear latent dimensions. Considering all algorithms together, coverage increased with latent dimensionality, but the amount contributed to the overall coverage by each model varied substantially (**Fig. 2D**). β-VAE contributed the largest proportion of unique gene set coverage (38%), with β-TCVAE and Vanilla VAE also showing strong contributions. Linear models (PCA, ICA, NMF) provided smaller but non-negligible coverage, suggesting that their latent dimensions still capture and optimize distinct biological signals. Importantly, the stacked coverage curves demonstrated that while no single model, or grouping of models for one latent dimensionality, achieved full coverage. However, the complementary contributions across models and across dimensionalities approach high levels of saturation (87%; **Supplementary Figure 3**).

Together, these results highlight a trade-off between model stability, reconstruction accuracy, and biological interpretability. While some models, such as the β-VAE, offer fairly balanced performance across all metrics, others like the β-TCVAE may sacrifice stability for greater diversity in biological signals. In general, the results also show how applying the BioBombe multi-model approach dramatically increases and optimizes signature discovery compared to traditional single-model approaches.

### Interpreting latent dimensions with Gene Set Enrichment Analysis

To functionally interpret the latent dimensions, we applied GSEA to the weight matrices produced by each of the six BioBombe algorithms across the 28 latent dimensionalities. This analysis leverages curated gene set libraries, including Reactome [26] and CORUM [27]. GSEA connects these latent dimensions (i.e., latent embeddings) to well-defined biological pathways and protein complexes, enabling biological interpretations of gene process dependencies.

Across all models and dimensionalities, GSEA returned a total of 3,516,392 significant Reactome pathway associations based on the GSEA standards of False Discovery Rates below 5% and Normalized Enrichment Scores above 1 [25]. The results span 1,031 unique pathways (**Fig. 3A**). To assess the specificity of these results and rule out spurious enrichment, we conducted a control experiment in which we randomly shuffled weight matrices prior to GSEA. This exercise yielded only 126 total significant pathway associations, strongly supporting the biological relevance of the original, model-derived associations as successfully capturing structured and biologically grounded gene sets. We also compared BioBombe enrichment against a naïve baseline in which we applied GSEA directly to ranked gene dependency profiles for each cell line (**Supplementary Figure 4**). While both approaches recovered broadly overlapping Reactome programs, latent-derived enrichments identified a larger number of unique pathways (1031 vs. 894). Furthermore, among pathways identified by both methods, BioBobme latent representations produced stronger normalized enrichment scores in 76% of cases, indicating that latent dimensions concentrate biologically coherent signals beyond those captured by raw dependency rankings.

**Figure 3.**
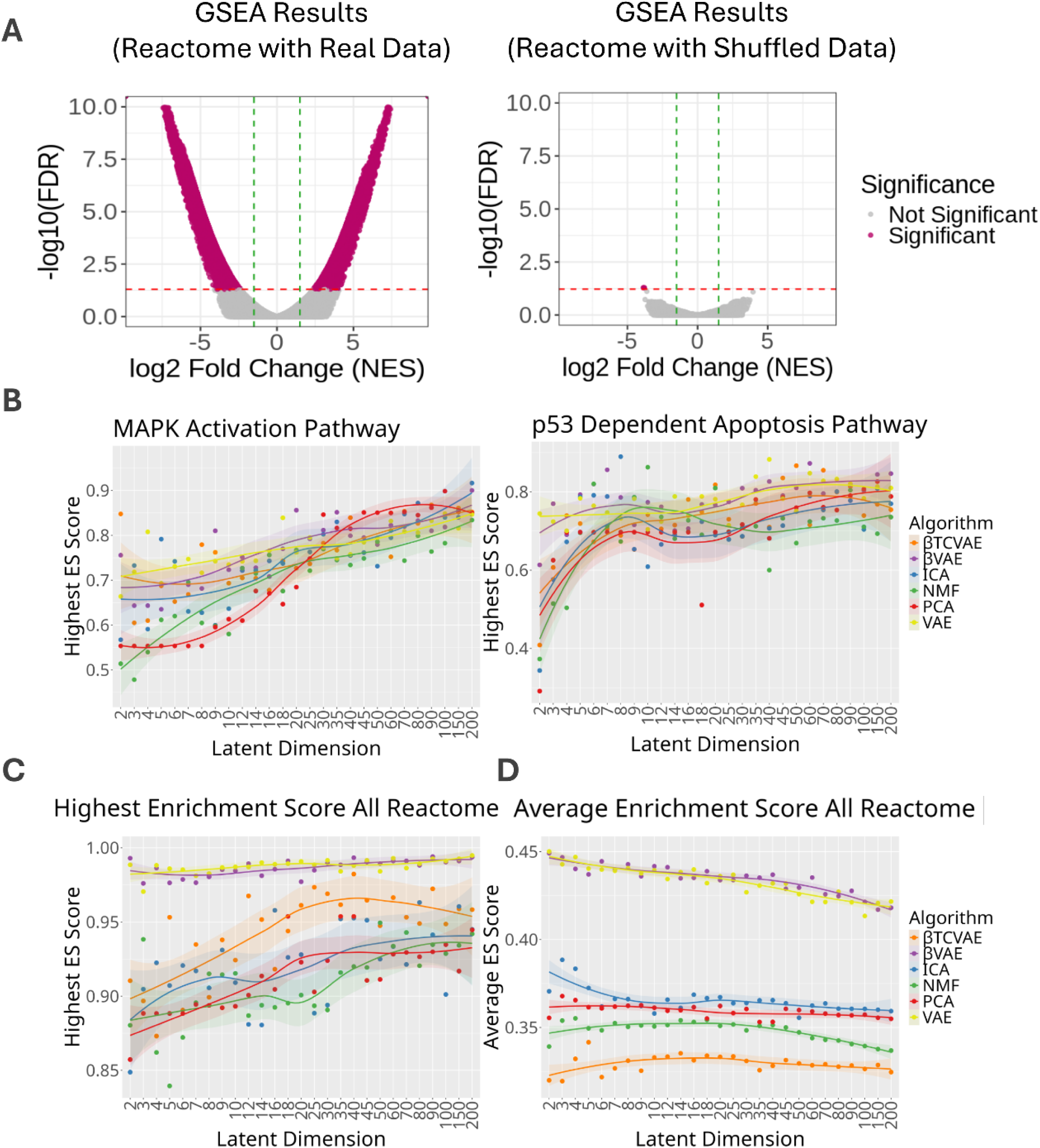
BioBombe optimizes gene process dependencies. **(A)** Gene Set Enrichment Analysis (GSEA) normalized enrichment score versus false discovery rate (FDR) for real (left) and shuffled (right) data applied to BioBombe weight matrices. **(B)** Comparison of models across latent dimensionalities using selected anecdotal Reactome pathways, showing the highest enrichment score (ES) per latent dimensionality. Comparison of models across latent dimensionalities, showing **(C)** highest overall enrichment and **(D)** average enrichment score for any pathway.

Biologically compelling pathways emerge across multiple BioBombe latent dimensions, which illustrate the strength of our approach. For example, we observe enrichment of the PINK1–PRKN mitophagy pathway, which can trigger cell death in response to mitochondrial damage, highlighting a known link between altered metabolic stress responses and tumor survival mechanisms [30]. In addition, BioBombe consistently implicates several FGFR-related signaling cascades. FGFR aberrations are recurrent in diverse cancer types and have been directly targeted by small-molecule inhibitors currently in clinical use [31]. Similarly, BioBombe consistently finds MAPK and WNT signaling pathways across many models, both of which govern cell proliferation and differentiation, and are well-established drivers of tumor initiation and progression [32, 33]. We also find recurrent enrichment of TP53-related pathways, underscoring the central dependency of p53 in DNA damage response, apoptosis, and resistance to therapy [34].

These enrichments not only demonstrate that latent dimensions capture fundamental gene process dependencies but also suggest clinically-actionable avenues, potentially grouping cancer samples by shared pathway vulnerabilities rather than single-gene effects. While the strength of enrichment varies across BioBombe dimensions and models, the consistent recovery of these pathways indicates that they represent robust, recurrent biological signals rather than model-specific artifacts. The distribution of latent scores further suggests heterogeneity in how strongly individual cell lines depend on these processes, raising the possibility of identifying tumor subsets particularly reliant on them. Furthermore, several of the implicated pathways, including FGFR and MAPK signaling, are the focus of current standard of care and ongoing drug development, directly connecting BioBombe-derived features to known therapeutic interventions.

### Optimizing gene process dependencies requires diverse models and latent dimensionalities

To understand how the BioBombe models optimize biologically relevant gene process dependencies, we further assessed how GSEA enrichment scores vary across latent dimensions and algorithms. Our results show that no single model or latent dimensionality universally outperforms the others, but distinct model-dimension combinations excel at capturing certain biological signals, which is consistent with prior BioBombe observations [14]. For example, the “*MAPK Activation*” pathway achieves its highest enrichment score with the β-TCVAE at *k* = 60 latent dimensions, while the “*p53 Dependent Apoptosis*” pathway is best captured by the β-VAE at *k* = 90 latent dimensions (**Fig. 3B**). These differences illustrate that even for well-characterized pathways, the optimal representation depends on both the architecture and the latent dimensionality of the model.

To move beyond focused examples of individual pathways, we next evaluated the full set of Reactome pathways to assess how well each model captured gene process dependencies across the entire library. In this broader analysis, we observed a general upward trend in maximum enrichment scores as the latent dimensionality increased, particularly for NMF and β-TCVAE, which benefited from the additional capacity to represent complex multi-gene patterns (**Fig. 3C**). Still, outliers such as PCA at *k* = 5 and ICA at low dimensionalities show that simpler or linear models can also efficiently optimize gene process dependency signals under the right modeling conditions. Complementing this analysis, we next examined the average enrichment score across all Reactome pathways (**Fig. 3D**). Unlike the peak enrichment trends in Figure 3C, which emphasize each model’s best-performing latent dimensions, the average enrichment analysis reflects the overall distribution of biologically-meaningful signals across the latent space. Here, the differences between algorithms were more dramatic, with VAE and β-VAE maintaining consistently higher average enrichment than all other methods. Interestingly, the β-TCVAE model that fell just behind the other VAEs in top enrichment scores falls to the bottom with averaged enrichment scores, showing that the high disentanglement penalties may allow that model to find several unique pathways quite highly, but it functions poorly globally.

### Correlating gene process dependencies with drug response

We next sought to evaluate whether the gene process dependencies identified by BioBombe are associated with drug response. To do this, we leveraged the PRISM drug repurposing dataset, which measures the viability of hundreds of matched cancer cell lines treated with thousands of compounds using a pooled barcoding approach [24]. We utilized the PRISM drug annotations because PRISM drug response measurements and DepMap gene dependency data are generated within the same experimental ecosystem, providing a consistent framework for correlating latent gene process dependencies with drug sensitivity. For each latent dimension, we calculated Pearson correlations between its sample-activation scores (z values) and the log-fold change values from PRISM. A strong positive correlation indicates a latent representation associated with increased resistance to a drug, whereas a strong negative correlation suggests the gene process confers drug sensitivity. Correlating drug responses with gene process dependencies enables both the discovery of novel drug targets and the repurposing of existing compounds for tumors with specific multi-gene vulnerabilities.

Across all models and dimensions, we identified 15,739 significant drug-latent correlations (**Fig. 4A**). To avoid counting the same association multiple times, we filtered results by unique drug correlations, selecting the highest correlating model. This process yielded 2,695 distinct significant correlations (511 unique drugs), which we will refer to as gene process dependency “hits.” In contrast, we randomly shuffled latent dataframes and observed only 37 significant associations (**Fig. 4A**). Additionally, we performed an analogous Pearson correlation analysis on the PRISM dataset using the naive GSEA enrichment scores, generated from the DepMap dataset without latent compression. This analysis yielded 155 unique significant drug associations. Comparing those drugs between methods revealed that 90.16% exhibited stronger absolute Pearson correlations in the latent-space analysis than in the naive gene set approach (**Supplementary Figure 5**), suggesting that the BioBombe-derived latent representations capture biologically relevant drug-response dependencies more effectively than more naive approaches.

**Figure 4.**
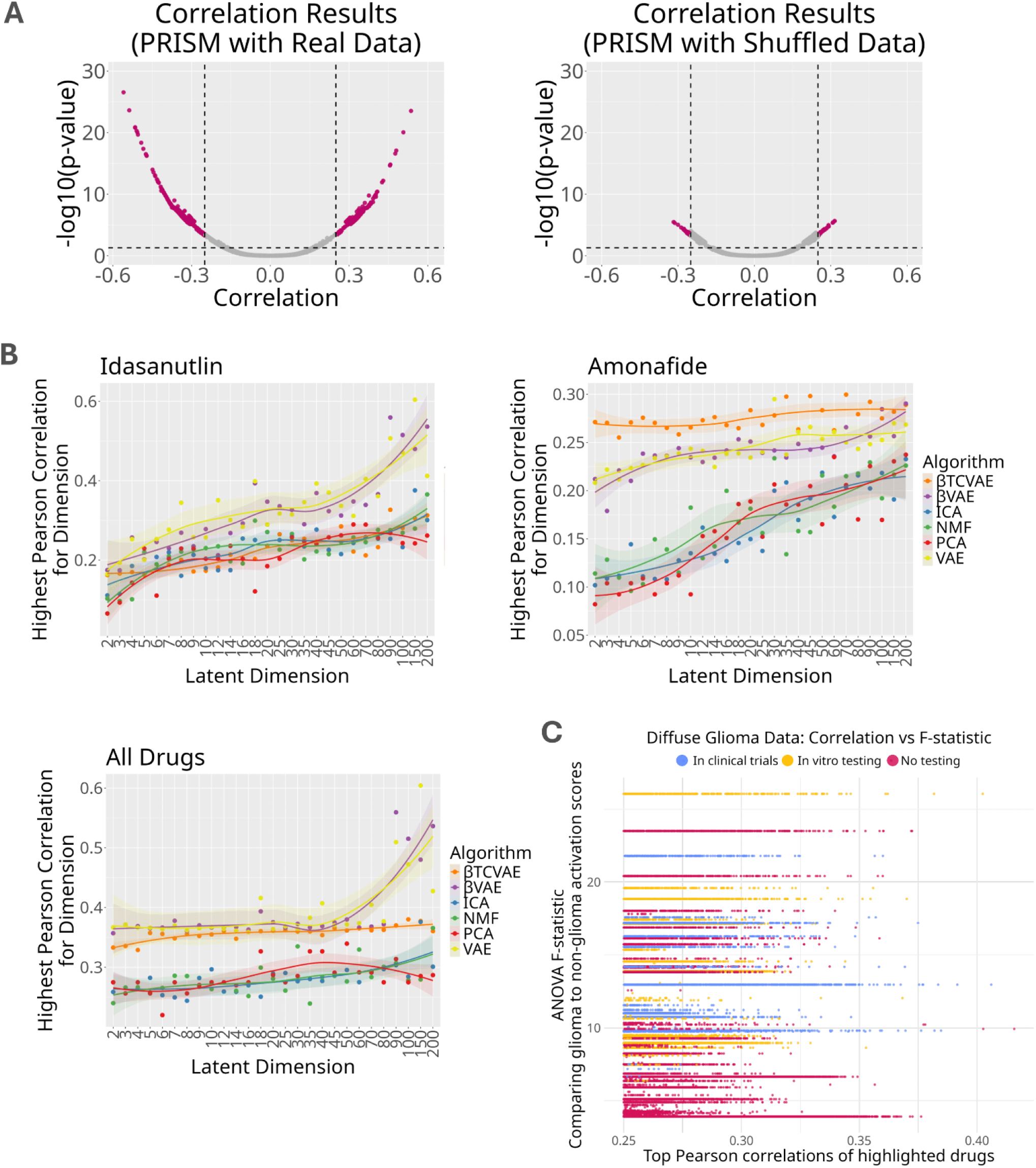
Correlating drug activity across BioBombe latent dimensions. **(A)** Correlation between BioBombe latent scores and PRISM scores. Significance cutoffs for correlation (+/-0.25) and p-value (0.05). **(B)** Correlation of selected anecdotal drugs (and all drugs) across *k* latent dimensions and models. **(C)** Glioma-specific drug analysis classifying drugs identified for glioma cell lines by clinical trial status: clinically tested, tested in vitro only, or untested. Comparing Pearson correlations of drug sensitivity and ANOVA F-statistics highlight drugs with significantly enriched activity in gliomas compared to all other cancer types.

To further characterize how model architecture and latent dimensionality influenced drug correlations, we quantified both the proportion of unique significant drug associations recovered by each model and the cumulative non-overlapping drug coverage across latent dimensionalities (**Supplementary Figure 6**). Consistent with the drug correlation analyses, the VAE-based models recovered substantially more unique drug associations than PCA, ICA, or NMF across nearly every latent dimensionality. Although increasing latent dimensionality initially expanded the number of recoverable drug associations, the rate of new drug correlations gradually diminished, with coverage approaching a plateau at higher dimensionalities. This contrasts with gene coverage analyses, where increasing latent dimensionality continued to reveal additional biological processes. These findings suggest that many therapeutically relevant gene process dependencies are represented by relatively broad latent features, whereas progressively higher-dimensional models primarily capture increasingly specialized biological variation that contributes comparatively fewer novel drug associations.

The top-correlated drug was idasanutlin, an MDM2 inhibitor [35], which displayed the highest correlation with latent dimension *k* = 150 from the VAE model (Pearson’s R = 0.61). This finding is consistent with the original PRISM study, which identified idasanutlin and other MDM2 inhibitors as top hits most closely aligned with CRISPR knockout of their molecular targets [24]. Notably, the PRISM study also demonstrated that such target-level concordance is relatively uncommon across the broader compound landscape, particularly among repurposed non-oncology agents, where only a small fraction of active compounds exhibited response patterns consistent with perturbation of their annotated targets. These observations suggest that drug sensitivity is frequently influenced by broader pathway-and network-level dependencies rather than direct target effects alone, supporting our use of latent gene process representations to capture higher-order biological relationships associated with therapeutic response. Additionally, amonafide, a DNA intercalating agent [36], exhibited a moderate correlation (Pearson’s R = 0.3) with a latent dimension characterized by unusually high activation scores across nearly all tumor samples (the highest average sample activation score (z-score) of any dimension) making it a distinct and biologically interesting case. Unlike pathway enrichment, where higher latent dimensionality often led to improved signal detection, drug-latent correlations exhibited more stable or plateauing behavior across dimensionalities, particularly in β-VAE and VAE, suggesting that drug-response relationships may be captured with fewer latent dimensions, or have less biological complexity (**Fig. 4B**). As a resource for future discovery, we present the significant drug-latent correlations and their associated gene process annotations in **Supplementary Table 1**.

To investigate whether BioBombe can uncover both known and novel therapeutic opportunities in specific cancer types, we performed a glioma-focused analysis by comparing drug activity in glioma cell lines to all other cancers. Diffuse gliomas, especially pediatric forms, remain among the most treatment-resistant cancers, with few targeted therapies and limited translational insights from adult studies [37]. Considering only latent dimensions that were significantly correlated with the activity of any drug, we calculated differential activity in glioma compared to non-glioma samples using an ANOVA test for each drug’s activity (**Fig. 4C**). We observed several clinically-approved glioma therapies, such as osimertinib [38] and gefitinib [39], showing both strong Pearson correlations with latent dimensions and high ANOVA F-statistics, indicating high, differential activity in gliomas. Likewise, drugs previously shown to be effective *in vitro*, such as ibrutinib [40], also emerged as candidates. However, BioBombe also surfaced an interesting set of untested drugs such as Ro-4987655, EVP4593, and BAY-87-2243 that show both glioma-specific differential activity and high correlations with highly-active gene process dependencies in gliomas. These candidates warrant further exploration, as their consistent associations suggest that they may exploit glioma-specific vulnerabilities not previously targeted in clinical or preclinical settings.

### Single-sample gene process dependencies and correlated drugs

To understand how multi-gene process dependencies manifest at the level of individual cell lines, we performed single-sample analyses across all DepMap. For each sample, we assigned a gene set to each latent dimension by associating it with the pathway, CORUM complex, or drug exhibiting the highest enrichment score (highest GSEA for pathway/complex and Pearson correlation for drugs). This framework allows us to transform each tumor’s latent profile into interpretable biological and pharmacological features, effectively generating a latent “pinwheel” of gene-process dependencies and candidate drugs for every sample (**Figure 5**). Note that this process is different from the dataset-wide annotation approach, which assigns pathways/complexes/drugs based on all samples (see Methods for more details). Each line on the pinwheel corresponds to a latent score from one of the selected dimensions, annotated with the highest-ranking associated gene process (i.e., Reactome pathway or CORUM complex) or drug. We present the top 15 drugs for each cell line in **Supplementary Table 2**.

**Figure 5.**
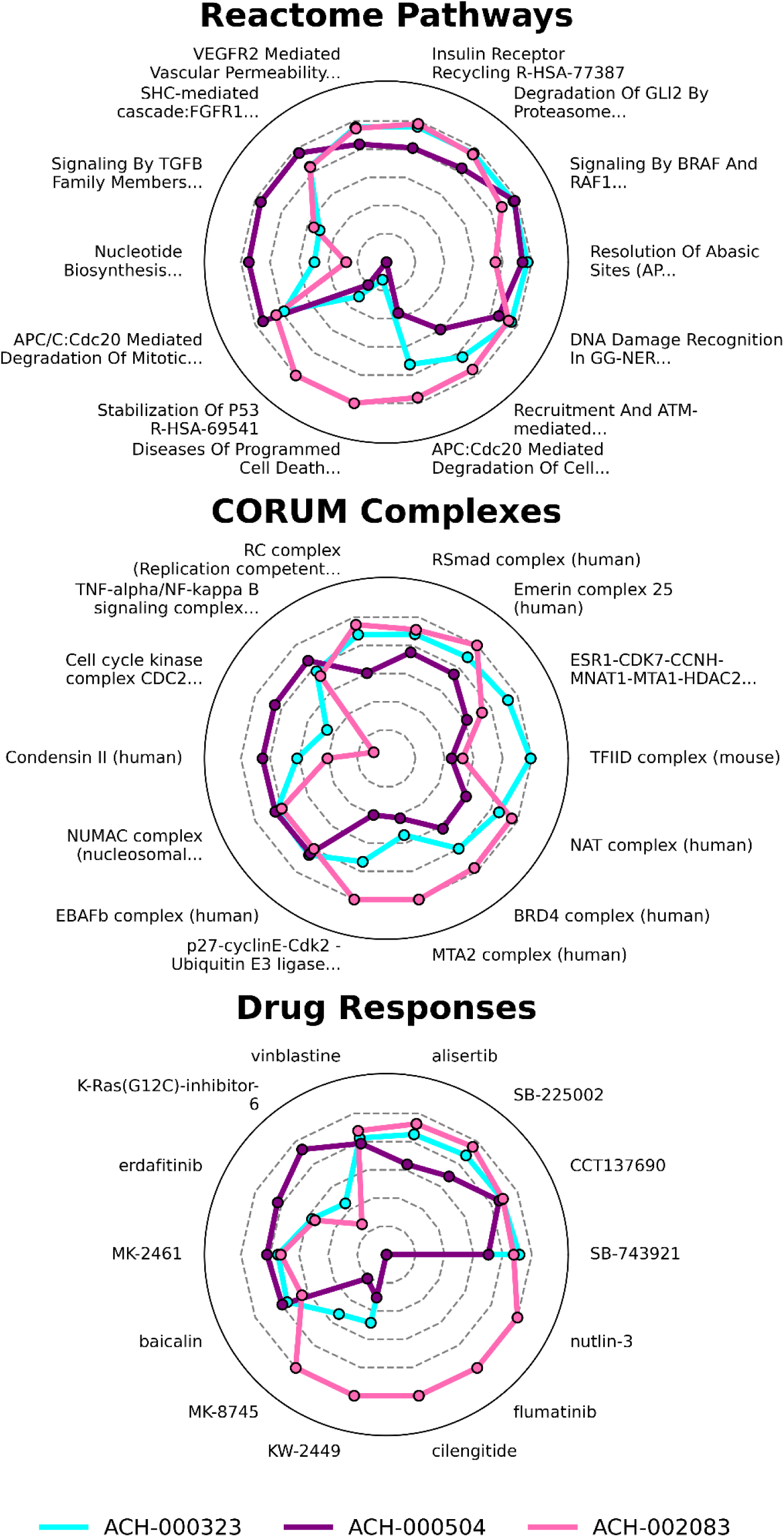
Single-sample latent dependency prediction pinwheels. Each color represents one tumor sample from DepMap, with Reactome pathway (top), CORUM complex (center), and drug (bottom) predictions derived from normalized BioBombe scores. We assign each latent dimension its top associated feature based on enrichment or correlation (without replacement), and the magnitude of each point is equal to the normalized latent score for that tumor. The normalized latent score is scaled from 0 to 1 within the single latent dimension.

**Figure 5** showcases three representative tumor samples. The cyan color shows ACH-000323, a diffuse glioma with a rich and diverse profile of top-ranked drug predictions. Notably, its profile includes both commonly-predicted drugs like SB-743921, which is a top hit in 42% of all DepMap cell lines, as well as more context-specific drugs like CCT137690, which is predominantly enriched in diffuse glioma samples (ANOVA F-statistic = 4.339). In addition, unique compounds such as flumatinib and cilengitide emerge as top-ranked candidates for this sample. While amonafide and cladribine show modestly negative average fold changes, alisertib stands out with a PRISM average fold change of-0.262 for brain tumors and a fold change of-2.66 in ACH-000323, which is the second most negative response of any brain sample recorded for that drug. This suggests BioBombe has the ability to identify sample-specific drug sensitivities based on the calculated gene process dependencies.

ACH-002083, a neuroblastoma sample shown in pink, demonstrates how this approach is supported by previously established pathway and drug knowledge. One of the top drugs for this sample is nutlin-3, an MDM2 inhibitor, while two top Reactome pathways are “Stabilization of p53” and “Diseases of Programmed Cell Death.” Although this sample is not included in the PRISM dataset, DepMap data confirm that ACH-002083 does not carry a TP53 mutation and has a relatively low mutational burden, further supporting the potential efficacy of MDM2 inhibition in this context. This also highlights the ability of BioBombe to make drug efficacy predictions in samples not profiled with the PRISM assay. In contrast, ACH-000504, a second diffuse glioma sample (shown in purple) exhibits a less diverse set of top drug hits, and none of the latent scores for these hits are close to 1. A known p53 loss-of-function mutation exists for this sample, and the PRISM data shows none of the MDM2 inhibitors are effective. Likewise, the latent scores are extremely low for the p53 pathways and related drugs that were shown highly in ACH-002083.

### Validating BioBombe-prioritized compounds in pediatric high-grade glioma cell lines

Pediatric high-grade gliomas (pHGGs), including diffuse midline gliomas (DMGs), remain among the deadliest childhood cancers. Few effective targeted therapies exist, and the rarity of patient-derived models for some pHGG subtypes limits opportunities for large-scale functional screening [41].

Genome-wide CRISPR dependency and drug screening datasets such as DepMap provide a comprehensive map of cancer vulnerabilities, but equivalent functional datasets do not exist for most primary pediatric tumors. To bridge this gap, we sought to predict gene process dependencies directly from tumor RNA-seq data, prioritize drugs using drug-latent correlations, and validate these prioritizations in targeted viability experiments.

We first trained ElasticNet regression models to predict each latent gene process dependency (i.e., each latent z representation) from matched DepMap RNA-seq data. Because DepMap contains paired transcriptomic and gene dependency profiles for the same cell lines, these models learned the relationship between gene expression and the latent dependency landscape identified by BioBombe. We optimized a separate model for each of the 20,862 latent dimensions, evaluating each gene process dependency independently based on its own predictive characteristics. Performance varied substantially across latent dimensions. For example, the best-performing model achieved an R^2^ of 0.8817 predicting latent dimension 75 in a VAE trained with k = 150 total dimensions (**Supplementary Figure 7**), and the second-best achieved an R^2^ of 0.8815 predicting latent dimension 0 in a β-VAE trained with k = 80 total dimensions. We retained only latent dimensions with R^2^ > 0.4, a threshold reflecting moderate predictive performance that reduced the likelihood of propagating poorly predicted dependencies into downstream drug prioritization.

We next applied these trained models to RNA-seq data from pHGG patient-derived samples collected through Children’s Hospital Colorado [42], generating predicted latent dependency scores directly from each tumor’s RNAseq data. Comparing these scores with the previously identified latent drug associations allowed us to prioritize candidate therapeutics without requiring functional screening of the patient samples themselves. From these predictions, we selected three compounds for experimental validation (axitinib, cladribine, and 3-deazaneplanocin A) based on their strong latent-drug associations, mechanistic relevance to the predicted biological processes, and reported ability to penetrate the blood-brain barrier, which is essential for targeting brain tumors (see **Methods** for details).

For the BioBombe-prioritized drugs and selected cell lines, we applied a standardized MTS dose-response assay protocol to assess cell viability and determine IC50 values for each drug (**Supplementary Figure 8**). The MTS dose-response assays revealed differential sensitivity of pediatric glioma cell lines to the three BioBombe-prioritized compounds (**Supplementary Figures 9-11**; **Supplementary Table 3**). Axitinib produced measurable inhibitory responses across nearly all evaluated cell lines, with IC50 values spanning the low nanomolar to submicromolar range. Cladribine showed more heterogeneous responses, with several cell lines exhibiting IC50 values above the highest concentration tested. 3-deazaneplanocin A generally required higher concentrations to achieve comparable growth inhibition, indicating lower overall potency under the conditions tested. We next compared these experimentally measured sensitivities with the latent dependency scores predicted from each cell line’s transcriptome (**Figure 6**). Cladribine showed the strongest correspondence between predicted and measured sensitivity (R^2^ = 0.5), despite its heterogeneous viability responses.

**Figure 6.**
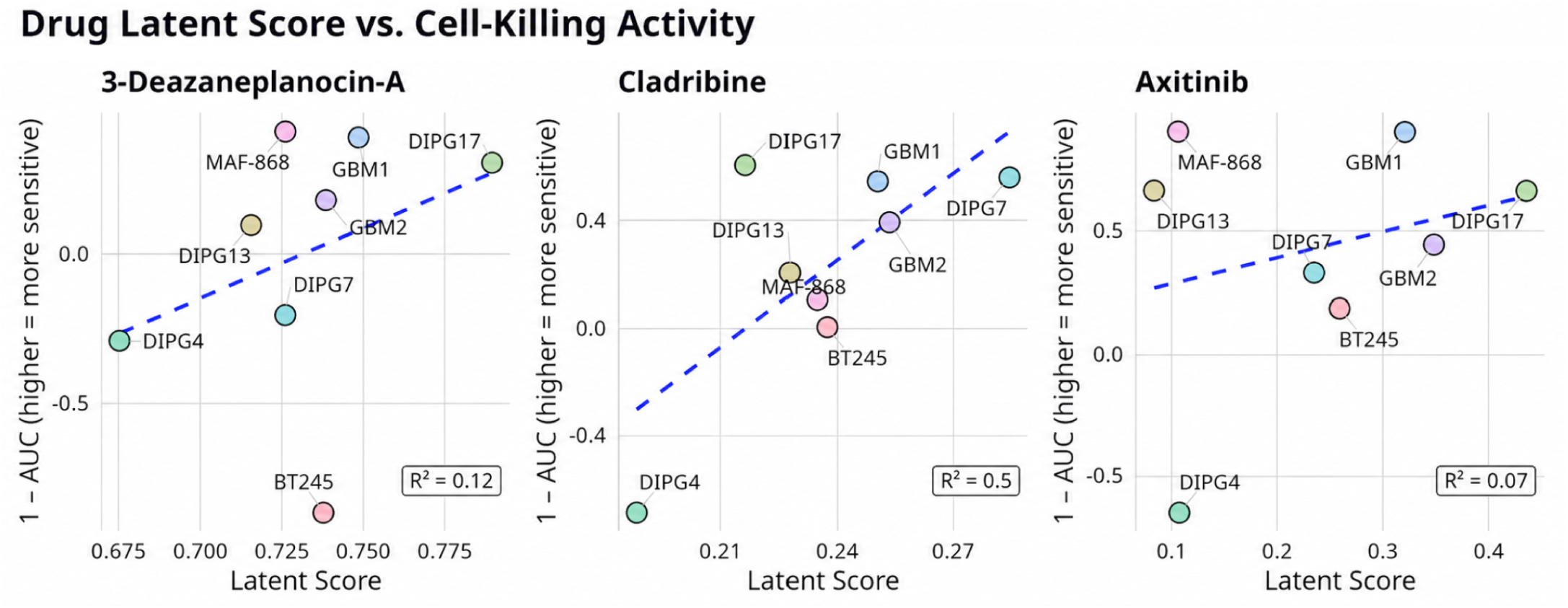
Pediatric high-grade glioma cell-killing validation. Each point represents one pediatric glioma sample.

3-deazaneplanocin A showed a weaker relationship between predicted and measured sensitivity, consistent with its lower potency overall. Axitinib showed the weakest correlation of the three compounds, though the relationship still followed a clear positive trend, suggesting the model captured a portion of the biological variation underlying response. Together, these findings provide proof of concept that latent gene process dependencies inferred solely from RNA-seq data can prioritize therapeutically relevant compounds in pHGG, while also highlighting the variability expected when translating predictions derived from large, heterogeneous cancer cell line collections to rare pediatric tumor models.

## Discussion

Applying BioBombe to gene dependency data introduces a powerful and scalable approach for uncovering complex gene process dependencies in cancer. By moving beyond single-gene dependencies, which has been a widely successful approach but that has historically limited the scope of therapeutic target discovery, we provide a method for identifying higher-order biological processes that underlie cancer vulnerabilities, which may better reflect actual drug activity given polypharmacology and off-target effects.

Our results demonstrate that different models and dimensionalities capture distinct biological signals. Both β-VAE and Vanilla VAEs consistently achieved strong performance in terms of reconstruction error and representation stability, as assessed by MSE and CKA, respectively. Notably, β-TCVAEs introduced greater variability, suggesting that the total correlation regularization drives the model toward uniquely structured, but less stable, latent dimensions. Future work could improve latent feature robustness through consensus modeling across multiple random initializations, stability-guided selection of latent dimensions, and incorporation of biologically informed constraints or weak supervision. Investigating ensemble or seed-averaged latent representations may further reduce stochastic variability while preserving biologically relevant structure and findings. This diversity of signal across models underscores the value of the ensemble approach used by BioBombe, suggesting that biological signal detection is not found best by one specific, optimized algorithm or dimensionality.

Different algorithms and latent dimensionalities may isolate separate aspects of the underlying gene process dependencies. Additionally, linear methods such as PCA offer improved interpretability and may be preferable when mechanistic clarity is the primary goal, whereas non-linear models such as β-VAE and β-TCVAE more effectively capture higher-order structure and broader biological programs. Notably, we observe that simpler models can occasionally recover pathway enrichments not identified by more complex architectures, highlighting that model complexity does not monotonically determine biological insight. As a result, there is no single “best” model or dimension. Instead, the optimal strategy is to embrace model and dimension diversity.

Gene Set Enrichment Analysis (GSEA) linked many latent dimensions to known biological pathways and protein complexes from Reactome and CORUM, including key processes like TP53 signaling, the citric acid cycle, and mitotic checkpoints. These findings highlight the biological interpretability of the latent dimensions, validating that our models are capturing real, biologically relevant gene combinations. The integration of PRISM drug screening data further revealed that certain gene process dependencies are significantly correlated with drug efficacy across cancer cell lines. For example, in gliomas, latent dimensions associated with TP53 and mitochondrial function were strongly correlated with the efficacy of drugs like osimertinib and gefitinib, as well as novel candidates like Ro-4987655 and EVP4593, supporting the potential for drug repurposing based on learned gene process signatures.

To explore the translational potential of these predictions, we investigated representative tumor samples with distinct drug prediction profiles to determine whether the prioritized compounds aligned with known cancer vulnerabilities and sample-specific biology. Several of the top-ranked candidates highlighted by BioBombe have mechanistic features that support their potential relevance in diffuse glioma. For example, ACH-000323, a diffuse glioma sample with a diverse set of predicted drug sensitivities, identified both broadly recurrent compounds such as SB-743921 and more context-specific candidates including CCT137690, flumatinib, and cilengitide. While SB-743921 represents a frequently identified compound across DepMap cell lines, CCT137690 showed enrichment among diffuse glioma samples, suggesting that BioBombe can distinguish general dependencies from cancer-type-associated vulnerabilities. Additionally, alisertib demonstrated a particularly strong predicted and observed association in this sample, with one of the strongest PRISM responses among brain tumor samples for this compound, supporting the ability of latent gene process dependencies to prioritize sample-specific drug sensitivities. Other predicted candidates, including amonafide and cladribine, are DNA-intercalating [43] or antimetabolite [44] agents that target proliferative cells, aligning with the high-replicative phenotype of diffuse gliomas. Together, these findings suggest that BioBombe captures multiple classes of actionable gene process dependencies, ranging from broadly conserved cancer vulnerabilities to more individualized therapeutic opportunities.

Experimental validation confirmed that BioBombe-prioritized compounds have measurable biological activity in pHGG models, though the magnitude and consistency of response varied across drugs and cell lines. Axitinib showed the broadest and most consistent antiproliferative activity, yielding quantifiable IC50 values in nearly all models tested, with low nanomolar potency in several. Cladribine also showed substantial activity but more heterogeneous responses: several cell lines failed to reach 50% growth inhibition within the tested concentration range. Despite this variability, cladribine showed the strongest correlation between predicted latent dependency scores and measured drug sensitivity, suggesting the latent dependency framework captures biologically-meaningful determinants of response beyond potency alone. In 2023, Kang et al. identified cladribine, an FDA-approved drug, as a compound that activates DGKB (diacylglycerol kinase beta) and suppresses DGAT1 (diacylglycerol O-acyltransferase 1), two enzymes involved in lipid metabolism [45]. They found that activating these enzymes with cladribine sensitizes glioblastoma cells to radiotherapy. Our results corroborates cladribine as a potential effective therapy with cell killing properties. In contrast, 3-deazaneplanocin A inhibited proliferation in multiple models but generally required higher concentrations for comparable effect, and showed weaker agreement between predicted and observed sensitivity. Notably, dose-response modeling in DIPG4 failed to yield stable IC50 estimates for any of the three compounds, consistent with the absence of a reproducible sigmoidal response under these conditions. While not every predicted association produced a strong quantitative correlation with drug response, identifying multiple compounds with demonstrable activity in independent pediatric glioma models supports latent gene process dependencies as a framework for therapeutic prioritization in rare cancers where large-scale functional screening is not feasible

While promising, our approach has several limitations. The 24Q2 DepMap dataset includes 1,150 cell lines, but some cancers (especially rare pediatric cancers) are absent or represented by very few samples. After filtering out samples with missing metadata, our BioBombe models were trained on 653 samples. Given our results, together with the precedent of the original BioBombe study training on one consortium with 734 samples, this sample size appears sufficient (see **Methods** for more details).

Reprocessing data as it becomes available will likely improve BioBombe’s performance. For example, integrating the growing influx of data from the Pediatric Cancer Dependencies Accelerator will likely increase multi-gene dependency discovery in rare pediatric cancers, while applying BioBombe to next-generation 3D cancer models may reveal additional biological signals [46]. Variability in CKA across latent dimensions indicates that not all latent dimensions and models are equally robust or interpretable, and many models probably aren’t learning biologically-meaningful signals. This highlights the necessity of filtering only to latent dimensions with optimized GSEA scores or drug correlations for future downstream applications. Additionally, although our pipeline supports drug prioritization,

Pearson’s Correlations for drugs are generally lower than we expected. This is likely due, in part, to off-target effects, polypharmacology, and other non-specific drug activities that are not well represented in gene dependency data. Furthermore, drug response is influenced by additional factors such as pharmacokinetic and pharmacodynamic effects, compensatory signaling pathways, cell state, and experimental variability. As a result, the relationship between latent gene process dependencies and drug sensitivity is inherently more complex than the relationship between latent scores and pathway enrichment. Despite these challenges, we observed multiple biologically-plausible drug-gene process associations, suggesting that the approach can still generate useful therapeutic hypotheses.

Experimental validation remains necessary to confirm the therapeutic efficacy and clinical relevance of prioritized drug candidates. Furthermore, because drug target annotations were used primarily for interpretation rather than for calculating latent-drug correlations, differences among annotation resources were not evaluated in this study. Future work could investigate whether integrating alternative drug target databases improves interpretation of latent-drug associations. Furthermore, a subset of samples exhibited few or no significant latent scores. This observation may reflect limitations of the current models or dataset, but it may also indicate biologically-meaningful convergence across tumors, such as shared dependency states, high mutational burden, multi-drug resistance, or other molecular features not captured in the present analysis. Additional investigation will be required to distinguish between these possibilities.Our transcriptomic prediction framework relies on RNAseq as an indirect surrogate for functional gene dependency. While matched DepMap RNA-seq and CRISPR dependency data enabled us to train Elastic Net models that accurately predicted many latent dependency programs, gene expression alone cannot fully capture cellular dependency. Functional dependencies are also influenced by genomic alterations, epigenetic state, post-translational regulation, protein abundance, metabolic context, and interactions with the tumor microenvironment. Furthermore, only a subset of latent dimensions could be predicted with sufficient accuracy (R^2^ > 0.4), indicating that some biological processes are not readily inferable from transcriptomic data alone. Consequently, predictions generated from RNA-seq should be viewed as estimates of latent dependency states rather than direct measurements. Future efforts will focus on integrating additional data types (e.g., proteomics, epigenomics) and validating predictions experimentally, further advancing the utility of gene process dependency modeling in translational cancer research and precision oncology.

## Methods

### Data Sources and Preprocessing

We used two primary datasets from the Broad Institute’s Cancer Dependency Map (DepMap) project:

### DepMap Achilles Gene Dependency Data

We used the CRISPR knockout data from the Achilles project (DepMap Public Version 24Q2) as the foundational dataset for identifying gene process dependencies across a wide range of human cancer cell lines (1,150 total lines) and cancer types (73 total). We filtered cell lines that did not include age or cancer type metadata, which reduced our sample size to 933 cell lines. We used the DepMap AgeCategory column, which defined “pediatric” as below age 18 and “adult” as age 18 and above. These gene effect scores (corrected CERES scores) represent the degree to which a gene is essential for cell survival. We downloaded data directly from the DepMap portal (https://depmap.org/portal/download/) and filtered the original 18,443 genes to only those samples that passed the QC metrics defined by the Webster method [12], which filtered the genes based on high variance and perturbation confidence. We then additionally removed ribosomal gene families known to consistently oversaturate signals in latent representations, leaving us with 2,772 genes that passed both QC methods. We also standardized gene names, selecting the Hugo gene symbols to align with GSEA libraries.

### PRISM Drug Repurposing Data

We used drug sensitivity scores (log fold-change values) from the PRISM Repurposing dataset, which measures compound-induced changes in viability across hundreds of barcoded cancer cell lines screened in pooled format, to evaluate the potential association between gene process dependencies and pharmacological response of small molecule drugs. PRISM reports log2-fold changes in cell viability for each compound-cell line pair, providing cell line-specific measurements of drug sensitivity rather than aggregate responses across cell lines. We obtained PRISM data from the DepMap portal (24Q2 release) and filtered drugs to retain only drugs with non-null values across at least 100 cell lines. We included a total of 4686 compounds and 578 overlapping cell lines in our analysis. We aligned BioBombe latent scores and PRISM response profiles using DepMap ModelID identifiers, and we calculated Pearson correlations across the shared cell lines for each latent dimension-drug pair.

### BioBombe Framework

We extended the original BioBombe framework to construct a large-scale ensemble of unsupervised machine learning models applied across multiple latent dimensionalities. We implemented the following models: Principal Component Analysis (PCA) [17], Independent Component Analysis (ICA) [18], Non-negative Matrix Factorization (NMF) [19], Vanilla Variational Autoencoder (VAE) [21], Beta Variational Autoencoder (β-VAE) [22], and Beta Total Correlation VAE (β-TCVAE) [23].

PCA identifies orthogonal components that capture the greatest variance in the data, providing a linear baseline for comparison [17]. ICA decomposes the data into statistically independent components, potentially revealing non-overlapping biological processes [18]. NMF constrains all components to be non-negative, and learns by optimizing an iterative process which yields additive and interpretable features often aligned with biological activity patterns [19]. The vanilla VAE, a probabilistic nonlinear model, learns a latent representation that can capture complex, nonlinear dependencies across genes [21]. The β-VAE introduces a tunable regularization term that enhances disentanglement among latent features, promoting separation of independent biological signals [22]. Finally, the β-TCVAE explicitly penalizes total correlation to further encourage statistical independence of latent variables [23].

We chose these models to balance interpretability, diversity of learned representations, and capacity to capture nonlinear relationships. While the original BioBombe framework included a Denoising Autoencoder (DAE), we excluded it here due to consistently poor reconstruction performance, whereas the VAE demonstrated strong stability and biological interpretability across latent dimensions, so additional variants were included.

Furthermore, the original BioBombe framework was applied separately to 11,069 bulk RNA-seq samples from The Cancer Genome Atlas (TCGA) PanCanAtlas [47], 11,688 bulk RNA-seq samples from the Genotype-Tissue Expression project (GTEx) [48], and 734 bulk RNA-seq samples (seven tissues) from the Therapeutically Applicable Research to Generate Effective Treatments (TARGET) project [49]. Although applying BioBombe to RNA-seq data differs fundamentally from applying it to gene dependency data, these datasets spanned a comparably broad range of biological diversity (33 cancer types in TCGA, 53 tissue types in GTEx, and seven cancer types in TARGET; 73 in ours), and TARGET in particular was similar in sample size to this study’s dataset (n = 933).

### BioBombe Training

We set 70 percent of the data in the training set, 15 in validation and 15 in testing, balanced to include similar distributions of age and cancer types in each data split. We MinMax scaled each split from 0 to 1 independently. Our training set contained 653 samples. We trained each model at 28 different latent dimensionalities (k = 2, 3, 4, 5, 6, 7, 8, 9, 10, 12, 14, 16, 18, 20, 25, 30, 35, 40, 45, 50, 60, 70, 80, 90, 100, 125, 150, and 200), and we trained the VAE variants at five different random initializations. These were based upon the parameters used in the original BioBombe suite. With the six model types, we trained a total of 504 different compression models. To optimize the architecture and training parameters of VAE-based models, we incorporated Optuna [28], an automated hyperparameter optimization framework. For each model and dimensionality, we used to identify optimal values for key hyperparameters (where appropriate) including: Learning rate, Batch size, β (beta) for KL weight (β-VAE), γ (gamma) for total correlation weight (β-TCVAE), number and size of hidden layers. We optimized using a loss function including reconstruction error (MSE) and Kullback-Leibler divergence.

### BioBombe Evaluation

We evaluated model performance in three ways:

### Reconstruction Error

To evaluate model fidelity, we quantified reconstruction error as the mean squared error (MSE):

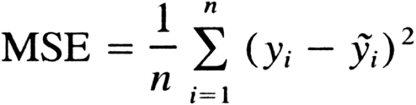

We calculate MSE between the input and reconstructed gene dependency matrices. For linear models (PCA, ICA, NMF), we calculated MSE reconstruction via the inverse transform of the latent representations. For neural network–based models (Vanilla VAE, β-VAE, β-TCVAE), we generated reconstructed outputs from the decoder, and computed MSE directly between the original and reconstructed matrices after inverse scaling to the original range. We averaged reconstruction metrics across test samples, with per-model results separated by latent dimensionality and initialization. To facilitate cross-model comparison, reconstruction errors were normalized by scaling between 0 and 1 using Scikit-learn MinMaxScaler within each model type before visualization.

### Latent representation stability

We evaluated stability and similarity using Centered Kernel Alignment (CKA) [29] to quantify similarity between learned feature spaces across model initializations, model types, and latent dimensionalities. CKA measures the degree of alignment between two sets of feature representations while being invariant to isotropic scaling, making it well-suited for comparing unsupervised embeddings that may differ in scale or orientation.

To compute CKA, we extracted weight matrices from trained models and computed centered Gram matrices for each representation. The similarity between two models X and Y was defined as:

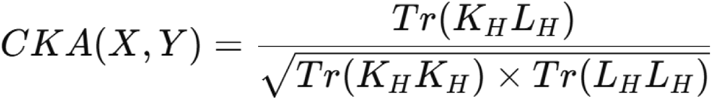

where K_H_ and L_H_ are the centered Gram matrices derived from X and Y, respectively.

For each latent dimensionality, we compared multiple random initializations of the same model type to assess within-model stability and averaged their CKA scores. Models included β-TCVAE, β-VAE, and Vanilla VAE, as these architectures are stochastic and sensitive to initialization. We then compared across-model stability by evaluating CKA scores between different model types trained at the same latent dimensionality (e.g., PCA vs. ICA, or NMF vs. β-VAE).

### Gene set coverage

Coverage quantifies the breadth of biological processes captured. We define gene set coverage as the total number of unique gene sets (from the Reactome and CORUM libraries independently) that are significantly enriched. To control for multiple hypothesis testing, we defined a significantly enriched gene set based on a Bonferroni correction (α = 0.05 / number of tested dimensions). We also considered gene sets with a Normalized Enrichment Score ≥ 1.5 (based on GSEA standards) to retain gene sets with high effect sizes. We evaluated coverage at three hierarchical levels to facilitate cross-model comparison:

1. Per-dimension–model coverage — the number of significant gene sets for a single latent dimension and model instance (specific model and initialization fit). To obtain model-level trends, we averaged the per-dimension coverage values across all initializations of the same model type, producing average coverage curves across latent dimensionalities. We also computed an overall (ensemble) coverage metric by taking the union of all significant gene sets across every model, initialization, and dimension. This value represents the fraction of all possible gene sets captured at least once anywhere in the ensemble.
2. Per-dimension coverage — the union of significant gene sets across all models and initializations for a single latent dimension. In a secondary, stricter analysis, we computed non-overlapping coverage, ensuring that each gene set was assigned to only one model per latent dimension to prevent double-counting of overlapping enrichments. This was done by sorting enrichments by GSEA enrichment score (|ES|) and retaining only the top gene set, found by a single model, for each specific dimensionality based on that score. This produced a non-overlapping model contribution plot showing how different model types contributed to overall biological coverage at a single dimensionality.
3. Ensemble coverage — the union of all significant gene sets across all models, dimensionalities, and initializations. We extended the strict non-overlapping approach across cumulative dimensionalities to evaluate how many dimensions are required for the ensemble to reach biological saturation. Coverage at dimension *k* includes all unique pathways discovered at lower dimensions, plus any newly discovered pathways at dimension *k*. This identifies the point at which additional latent dimensions no longer introduce new biological processes.

We then quantified coverage for each *(model, initialization, latent dimension)* triplet. We counted the number of unique significant gene sets, and divided by the total number of gene sets in the Reactome or CORUM library to obtain the coverage:

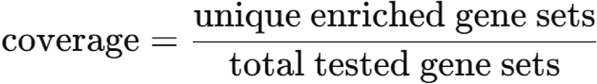

### Gene Set Enrichment Analysis (GSEA)

To interpret the biological relevance of each latent dimension, we applied Gene Set Enrichment Analysis (GSEA) to the model weight matrices using the BlitzGSEA Python package (version 1.3.33) [50]. For each trained model, we rank genes by their corresponding loading coefficients (weight magnitude and sign) from each latent dimension. We perform enrichment analysis for each dimension against the Reactome 2022 gene set library obtained from Enrichr [51]. We also repeat the analysis using the CORUM protein complex database to assess enrichment at the protein complex level.

For each latent dimension, BlitzGSEA computes normalized enrichment scores (NES), nominal p-values, and Benjamini–Hochberg–corrected false discovery rates (FDR). We define enrichment calls as significant using a log_2_ fold-change (LFC) threshold of 0.584 (corresponding approximately to a 1.5-fold change in enrichment score) and FDR ≤ 0.05, consistent with GSEA recommendations [25].

To assess the robustness of enrichment signals and control for potential artifacts in the model weight distributions, we generate 100 negative-control weight matrices by independently permuting gene loadings within each latent dimension and re-run GSEA on the permuted data. We compare the distribution of NES values from the real versus permuted analyses to confirm that observed enrichments exceed the null expectation. For each latent dimension, we record the top enriched gene sets ranked by NES, along with the direction of enrichment (positive or negative), nominal p-value, NES, and FDR.

### Drug Association via Pearson’s Correlation

To identify potential pharmacological dependencies associated with BioBombe-derived gene process dependencies, we correlate latent scores with drug response profiles of matched cell lines from the PRISM Repurposing dataset (DepMap release 24Q2). PRISM reports log_2_ fold changes in cell viability following treatment with thousands of small-molecule compounds across overlapping cell lines with the DepMap genetic dependency dataset.

For each model and latent dimension, we filter the latent dataframes to retain only those cell lines also profiled in PRISM. We align both datasets by ModelID identifiers and exclude columns with zero variance to prevent spurious correlations. For each latent dimension-drug pair, we compute Pearson correlation coefficients (r) using the scipy.stats.pearsonr function, yielding both a correlation value and an associated p-value. We repeat this procedure across all latent dimensions and all compounds tested in PRISM, producing a comprehensive latent-drug association matrix.

To facilitate biological interpretation, we retrieve drug-level metadata (including compound name, mechanism of action (MOA), molecular target, clinical phase, and indication) from the PRISM treatment annotations. We annotate each latent dimension with its top enriched Reactome pathways, as determined by Gene Set Enrichment Analysis (GSEA), to link molecular processes with associated pharmacological responses.

To assess the robustness of latent-drug associations and control for potential artifacts in the latent score distributions, we generate negative-control latent matrices by independently permuting the latent dependency scores across cell lines within each dimension. We recompute Pearson correlations for each permuted matrix, producing a null distribution of association strengths. We compare the real versus permuted correlation distributions to confirm that observed associations exceed the null expectation.

We define significant correlations as those with an absolute Pearson r ≥ 0.25, a threshold selected to highlight reproducible associations while minimizing noise across high-dimensional data. We visualize the resulting associations using volcano plots showing correlation magnitude versus statistical significance and compare trends across latent dimensionalities to identify model-and k-specific drug dependencies.

### Drug coverage

Coverage here quantifies the breadth of drug associations captured by each model. We defined drug coverage as the number of unique drugs associated with significant correlations to latent dimensions. A drug was considered significantly associated with a latent dimension based on the significance threshold used for the drug–latent correlation analysis. To facilitate comparison across models and latent dimensionalities, we evaluated drug coverage at multiple hierarchical levels.

1. *Per-dimension–model coverage* — the number of unique drugs with significant associations for a single latent dimension and model instance (specific model and initialization fit). To characterize model-level trends, we averaged per-dimension coverage across all initializations of the same model type, producing drug coverage curves across latent dimensionalities. We also calculated cumulative coverage by taking the union of all unique significant drug associations identified across the latent dimensions of each model.
2. *Non-overlapping drug coverage* — to quantify the incremental contribution of each model while avoiding double-counting associations identified by multiple models, we calculated cumulative non-overlapping drug coverage across latent dimensionalities. Drug associations were assigned based on the model with the strongest significant correlation, such that each unique drug association contributed only once to the cumulative coverage. This allowed us to assess how model architecture and latent dimensionality contributed to the discovery of novel drug associations.

### Single-Sample Inference and Visualization

To explore personalized applications of BioBombe, we performed single-sample prediction and visualization for every tumor sample in the DepMap dataset. For each sample, we matched the normalized latent z-scores to the highest-ranking enriched Reactome pathway, CORUM complex, and PRISM drug (based on Pearson’s correlation or enrichment score) using a non-replacement assignment strategy that ensured each latent dimension was uniquely associated with one relationship. This procedure yielded interpretable, per-sample latent dependency profiles that could be visualized as spider plots (“pinwheels”), where each axis represents a latent dimension annotated by its top-ranked biological or pharmacological feature.

To generate these visualizations, we implemented a plotting pipeline that computes latent-to-feature mappings and draws each sample’s profile as a polar “pinwheel” or bar chart using matplotlib. Each radial spoke corresponds to the magnitude of a normalized latent score, and top features are annotated along the circumference with label wrapping and rotation optimized for readability. For the representative examples shown in Figure 5, the three cell lines ACH-000323 (diffuse glioma), ACH-002083 (neuroblastoma), and ACH-002228 (diffuse glioma) were visualized across Reactome, CORUM, and PRISM drug associations to demonstrate inter-and intra-line variation in predicted dependencies and drug sensitivities.

Enrichment and correlation thresholds were selected to balance statistical rigor with biological interpretability. Specifically, we retained only features with an absolute enrichment score (|ES|) > 0.584 or a Pearson correlation coefficient |r| > 0.25 and *p* < 0.05, thresholds corresponding approximately to one standard deviation from the mean association strength. These cutoffs were empirically determined based on background null distributions across all latent dimensions to ensure consistent stringency across feature types and prevent spurious associations.

### Pediatric glioma cell line validation experiments

#### RNAseq prediction framework

We trained elastic net regression models to predict gene process dependency scores, represented by the latent (z) dimensions, from the DepMap provided RNA-seq expression data. We separately optimized a model for each of the 20,862 latent dimensions, evaluating model performance independently for each gene process dependency. We retained latent dimensions with a test-set R^2^ > 0.4 for downstream analyses. The resulting models were then applied to RNA-seq data from pediatric high-grade glioma (pHGG) and diffuse midline glioma (DMG) samples to predict their gene process dependency profiles in the learned latent spaces. These predicted dependency profiles were subsequently correlated with drug response profiles to identify compounds predicted to target the gene processes represented by each latent dimension. Thus, this approach enabled drug-gene process associations identified from BioBombe and trained on the DepMap cell lines to be translated to individual samples based solely on their RNA-seq profiles.

#### Selection of cell lines for validation

We evaluated pediatric glioma cell lines BT245, MAF868, DIPG4, DIPG7, DIPG10, DIPG13, DIPG17, GBM1, and GBM2 for sensitivity to the BioBombe-prioritized compounds axitinib, cladribine, and 3-deazaneplanocin A using MTS-based cell viability assays. The cell line panel includes DMG and pHGG models representing both H3K27M-mutant and H3-wild-type backgrounds.

#### Preparation of drug stocks and dose-response dilution

We dissolved compounds in dimethyl sulfoxide (DMSO) to generate concentrated drug stocks, divided each stock into aliquots, and stored them at −20°C to minimize repeated freeze-thaw cycles. We prepared dose-response dilutions on a separate 96-well drug dilution plate using half-log serial dilutions: for the highest concentration, we added 2 µL of drug stock to 198 µL of the appropriate culture medium, and we generated successive concentrations by transferring 31.6 µL of the preceding dilution into 68.4 µL of culture medium, producing approximately 3.16-fold decreases in concentration between successive conditions. Because we subsequently transferred 20 µL of each dilution into 180 µL of cell suspension (a further 10-fold dilution) we calculated each working drug concentration to account for this final step, ensuring that we achieve intended nominal treatment concentrations in the assay wells. We tested each concentration in three technical replicate wells (20 µL per replicate), for a final assay volume of 200 µL.

#### Treatment of cells

For treatment, we transferred 20 µL of each drug dilution into wells containing 180 µL of cell suspension, for a nominal treatment volume of 200 µL per well, and tested each drug concentration in three technical replicates. We included vehicle-treated (DMSO) wells as controls and incubated cells with drugs for five days at 37°C and 5% CO₂.

#### Assessment of cell viability

We assessed cell viability on day 5 using the CellTiter 96® Aqueous One Solution Cell Proliferation Assay (MTS assay; Promega, G3580). We added 20 µL of MTS reagent to each well and incubated plates for 2-5 h, depending on the cell line, before measuring absorbance at 490 nm on a Synergy 2 microplate reader (BioTek). We calculated background absorbance from medium-only control wells and subtracted it from experimental measurements.

#### Dose-response curve generation

We generated dose-response curves in GraphPad Prism and fit them by nonlinear regression using a four-parameter logistic model with variable slope. We estimated half-maximal inhibitory concentrations (IC50) from the fitted curves and reported them in nanomolar (nM), presenting technical replicate measurements as mean ± SD. We designated IC50 values as unstable when a stable estimate could not be obtained from the fitted curve, and we reported values >10,000 nM when 50% inhibition was not achieved within the tested concentration range.

## Computational resources

We performed all computations in Python (version 3.12.8) using numpy (version 1.23.0), pandas (version 1.5.2), pytorch (version 2.4), and matplotlib (version 3.6.0). To train BioBombe, we used a computer with an AMD Ryzen 9 5900X 12-core CPU with 64 GB of DDR4 RAM. The full BioBombe suite ran in about 35 hours on this computer.

## Code Availability

All code used to implement BioBombe, perform dimensionality reduction, run GSEA, train regression models, and generate figures is available at: https://github.com/WayScience/gene-process-dependencies[52]

## Acknowledgements

We would like to thank Ethan Cohen, Rose Doss, and Mark Woodnorth for early contributions to the code base. We thank Dave Bunten, Michael J. Lippincott, Erik Serrano, Cameron Mattson, and Jenna Tomkinson for performing code review.

## Supplementary Tables

**Supplementary Table 1:** *Significant drug efficacy and BioBombe latent dimensionality correlations and their associated gene process annotations*.

**Supplementary Table 2:** *Top 15 drugs per DepMap sample correlated to BioBombe gene process dependencies*.

**Supplementary Table 3:** *Characteristics of the nine pediatric glioma cell lines selected for in vitro MTS dose-response assays.* We tested the cell lines on the BioBombe-prioritized compounds axitinib, cladribine, and 3-deazaneplanocin A. The panel includes diffuse midline glioma (DMG) and pediatric high-grade glioma (pHGG) models representing both H3K27M-mutant and H3-wild-type backgrounds.

