## Supplementary Figures for "Characterizing the landscape of gene process dependencies in cancer"

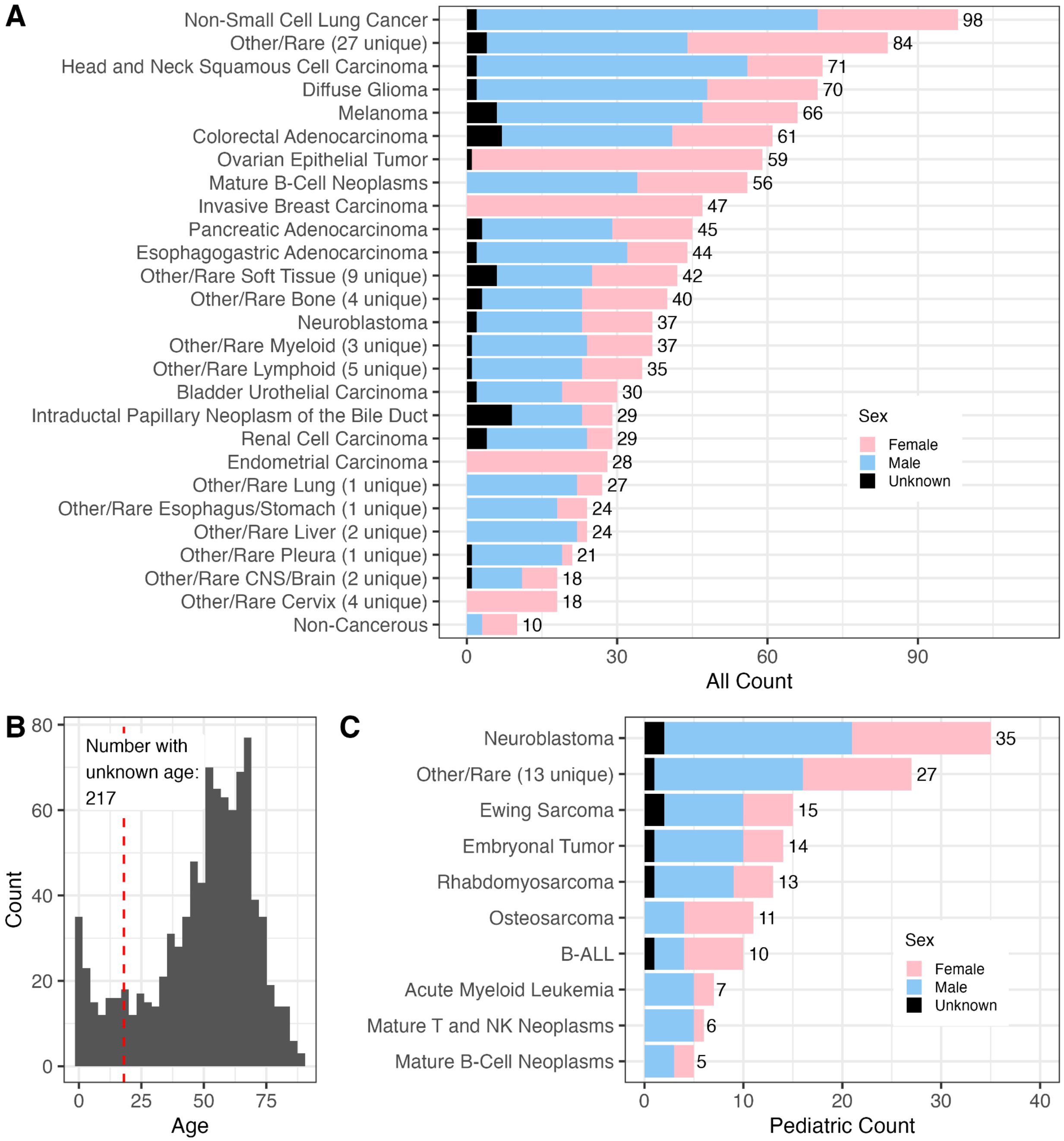

**Supplemental Figure 1:** *Cancer type and age distribution in 24Q2 DepMap gene dependency dataset.*

**(A)** Counts for all cancer types stratified by sex. There are 1,150 samples total. **(B)** Age distribution of cell line donors. The dashed red line indicates age 18, where we stratified adult and pediatric cancer cases. **(C)** Counts of pediatric cancer types.

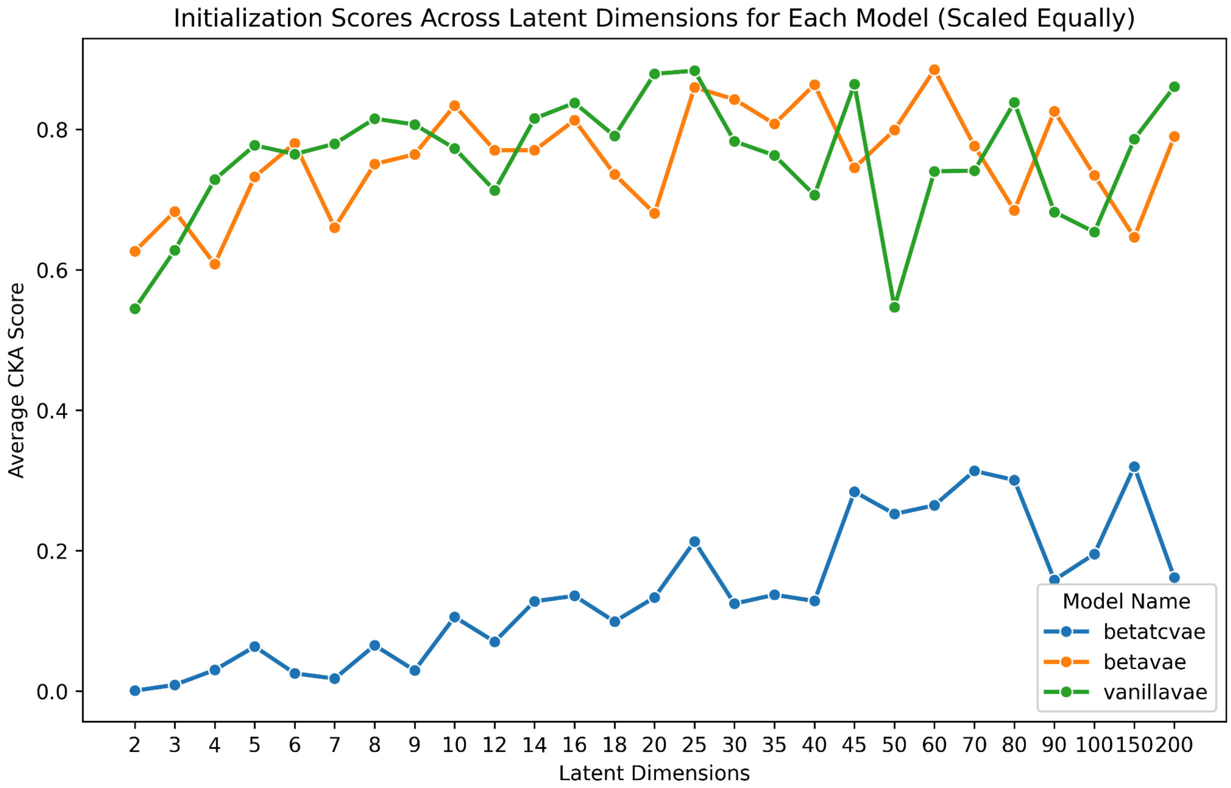

**Supplemental Figure 2:** *CKA scores for multiple initializations of VAE models*

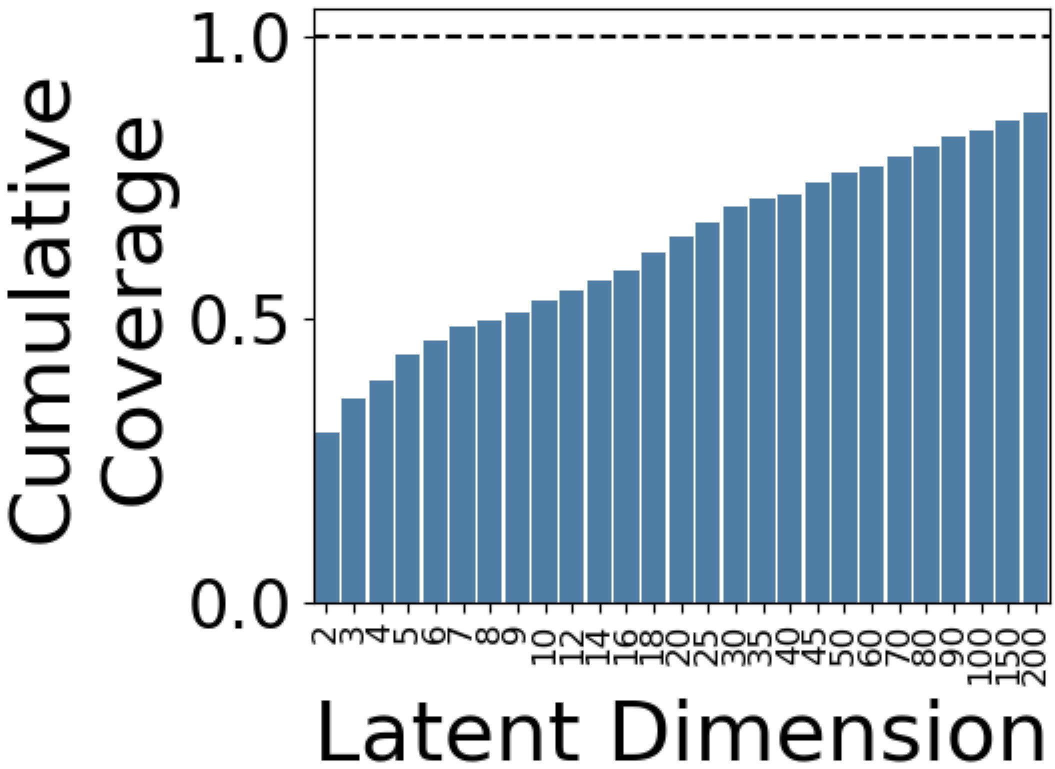

**Supplemental Figure 3:** *Ensemble Reactome gene set coverage, additive as dimensionality increases.*

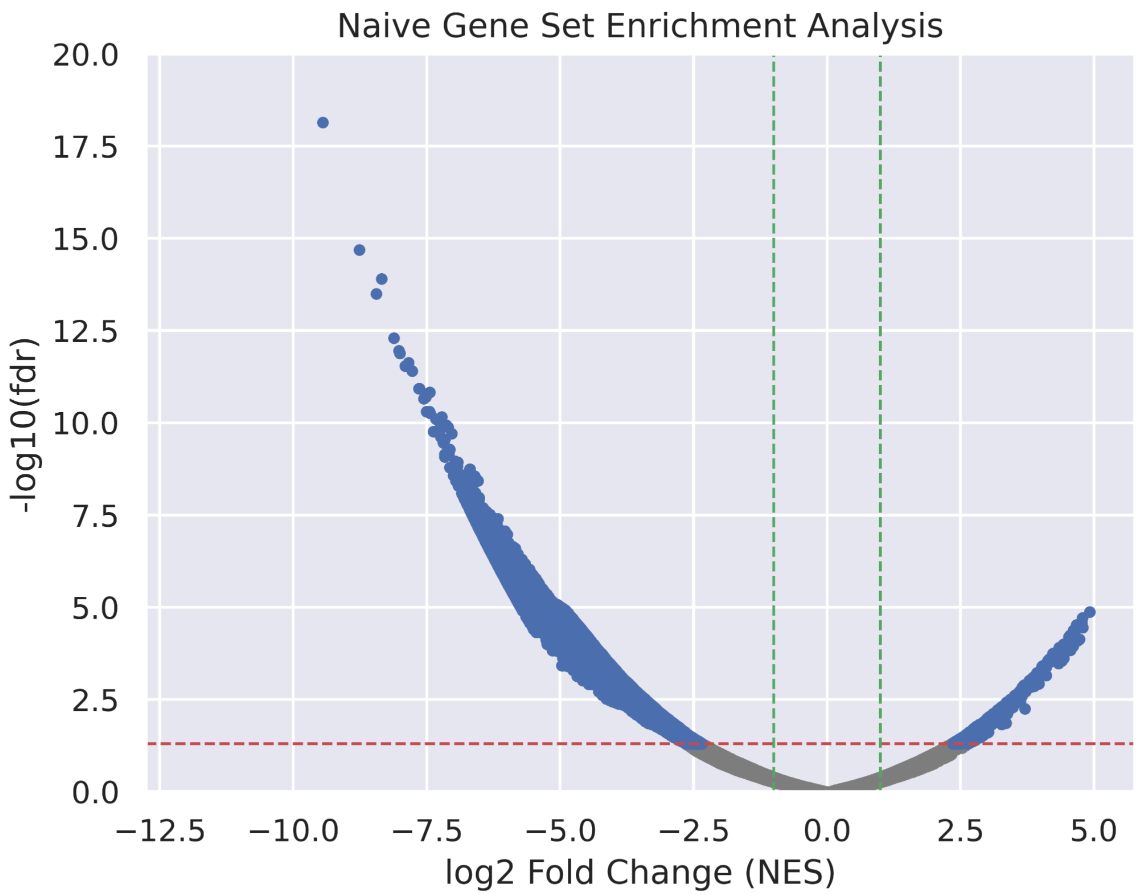

**Supplemental Figure 4:** *Naive Gene Set Enrichment Analysis, performed on the original, untransformed Achilles dataset.*

Normalized enrichment score (NES) versus false discovery rate (FDR). The red dotted line represents the significance cutoff for FDR, 0.05, and the green dotted line represents the significance cutoff for NES, 1.

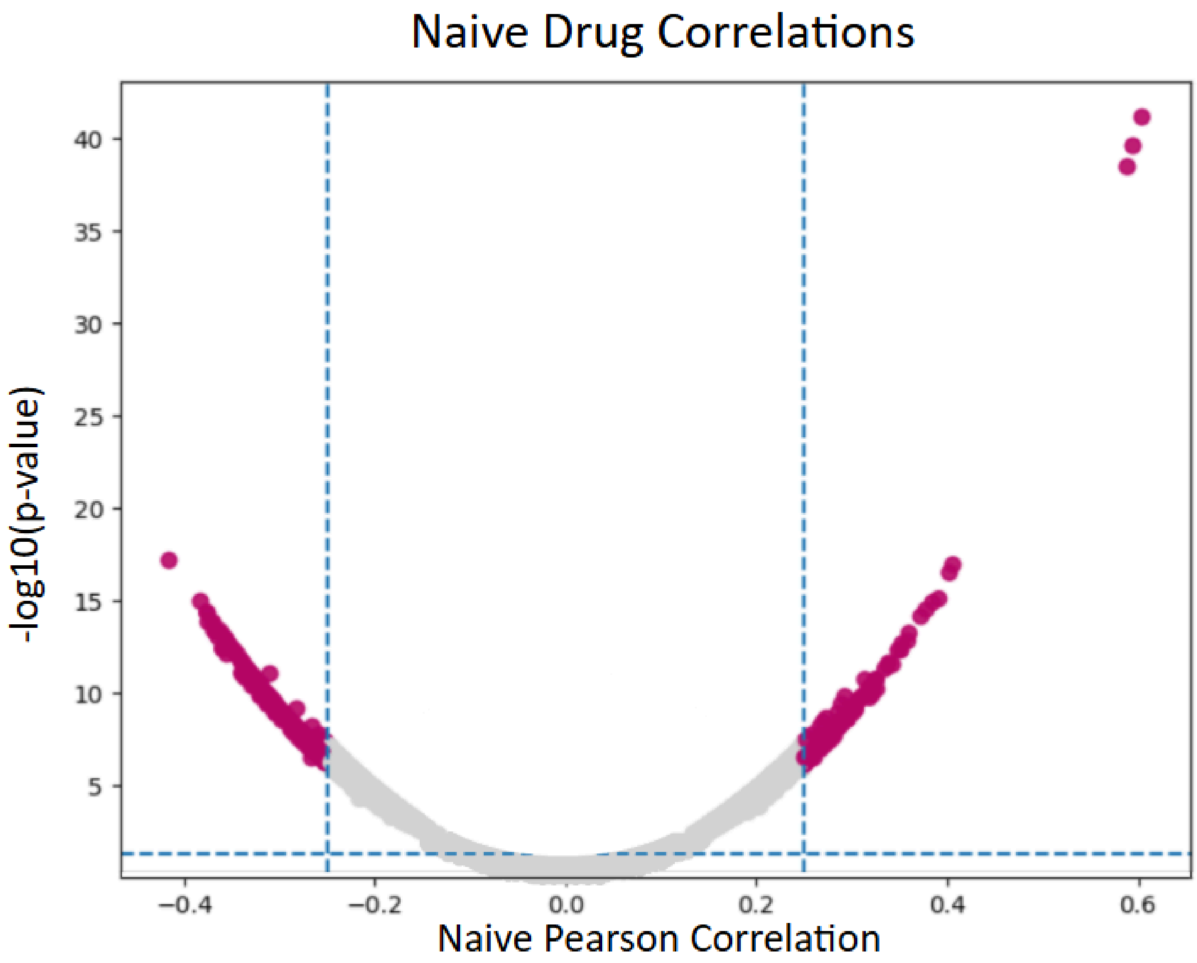

**Supplemental Figure 5:** *Naive drug correlations, performed on the DepMap dataset without BioBombe compression.*

We show Pearson correlation of DepMap gene dependency scores against PRISM drug activity. Dotted lines show significance cutoffs for p-value and Pearson correlation.

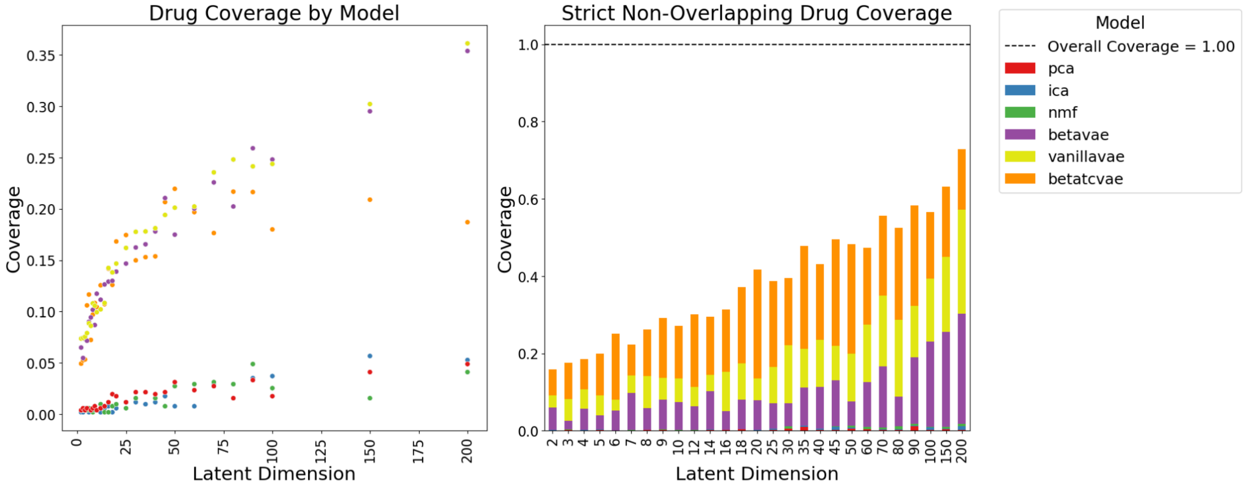

**Supplemental Figure 6:** *PRISM Drug Coverage.*

The proportion of total drugs recovered from the complete PRISM dataset, for each individual model/latent dimension combination and coverage for each dimension separated by model stacked for all models.

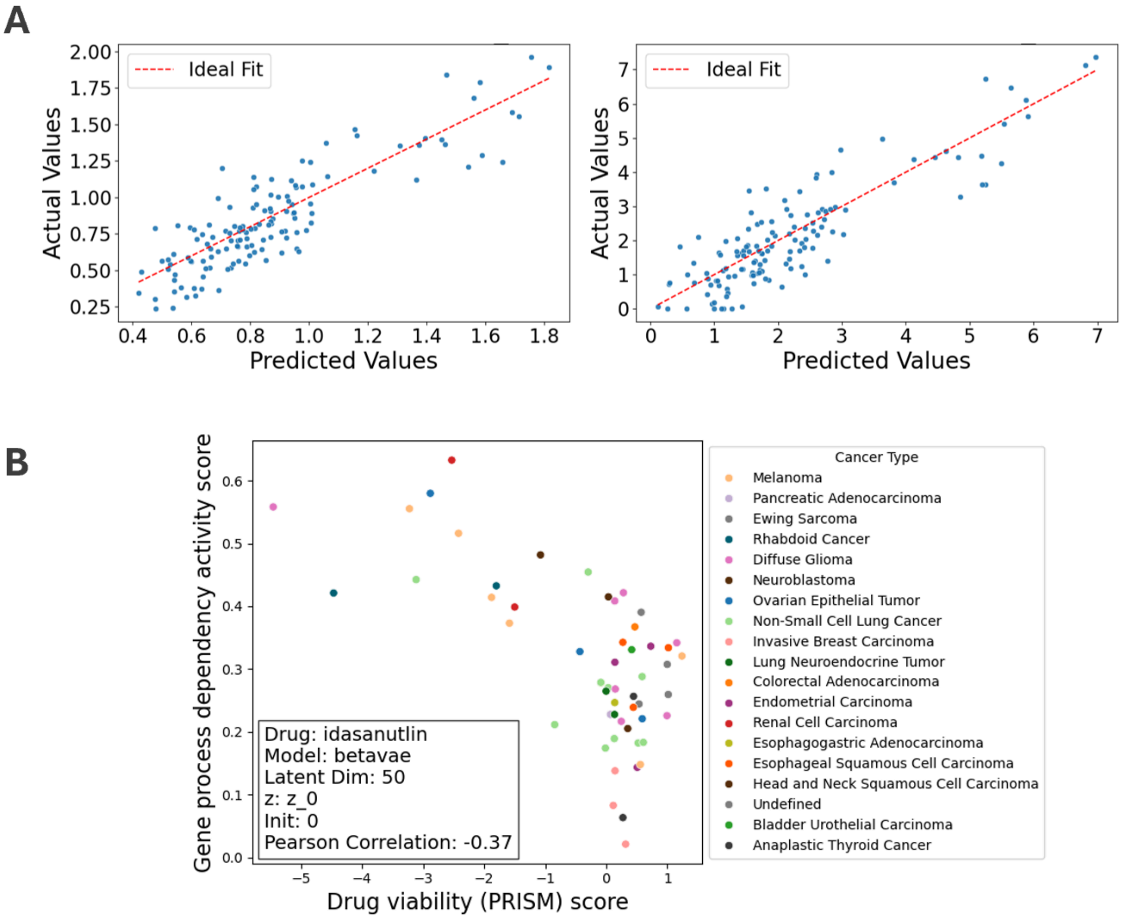

**Supplemental Figure 7.** *RNA-seq-based prediction of latent gene process dependencies and their application to drug sensitivity.*

**(A)** Predicted versus actual values for individual latent dimensions from elastic net regression models trained on DepMap RNA-seq data (two best performing models by $R^2$ shown). On the left is latent dimension z=75 in a VAE trained with k=150, and on the right is latent dimension z=0 in a β-VAE trained with k=80 total dimensions. Each point represents a single DepMap sample. **(B)** Predicted gene process dependency activity scores for idasanutlin vs corresponding PRISM drug viability scores across DepMap samples.

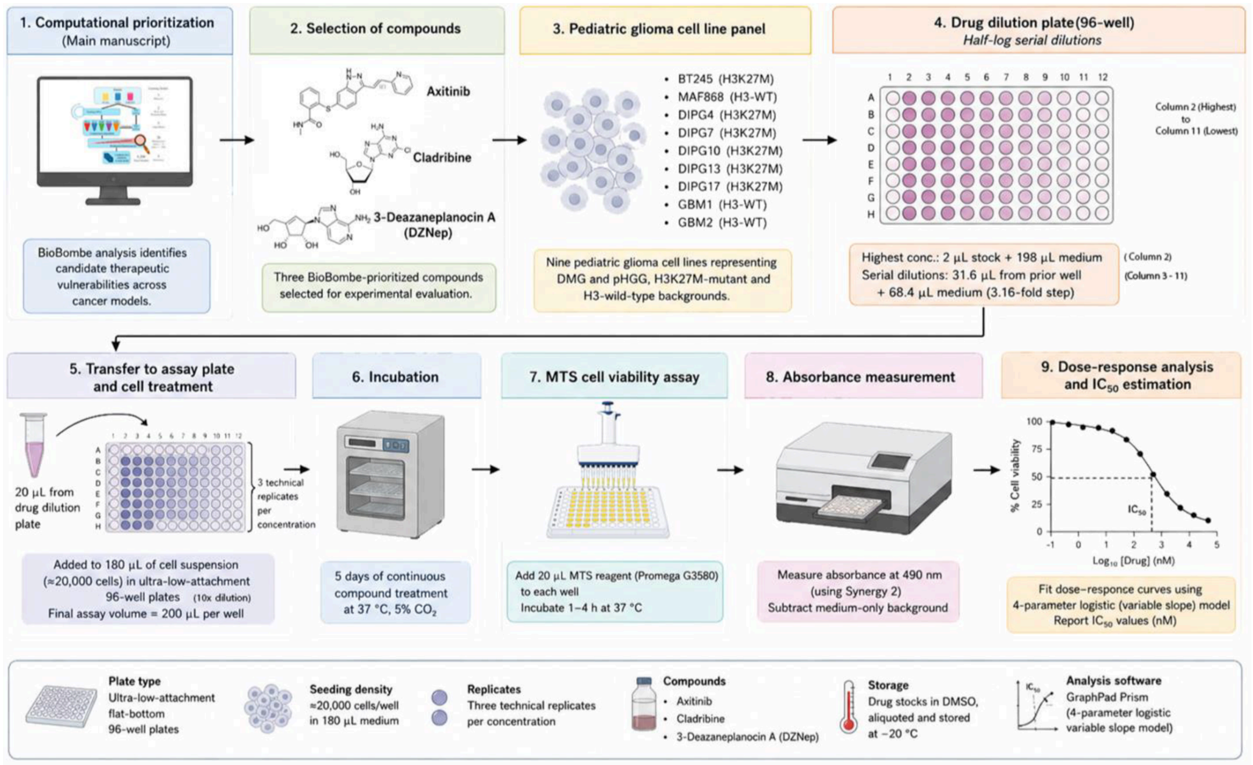

**Supplementary Figure 8:** *Experimental workflow for in vitro drug-response assays.*

We evaluated the BioBombe-prioritized compounds axitinib, cladribine, and 3-deazaneplanocin A (DZNep) across nine pediatric glioma cell lines. We prepared half-log drug dilutions on a separate 96-well plate and transferred them to cell-containing assay plates in three technical replicates per concentration. After five days of treatment, we assessed cell viability using the CellTiter 96® Aqueous One Solution Cell Proliferation Assay. We measured absorbance at 490 nm and fit dose-response curves by four-parameter logistic nonlinear regression in GraphPad Prism to estimate IC50 values.

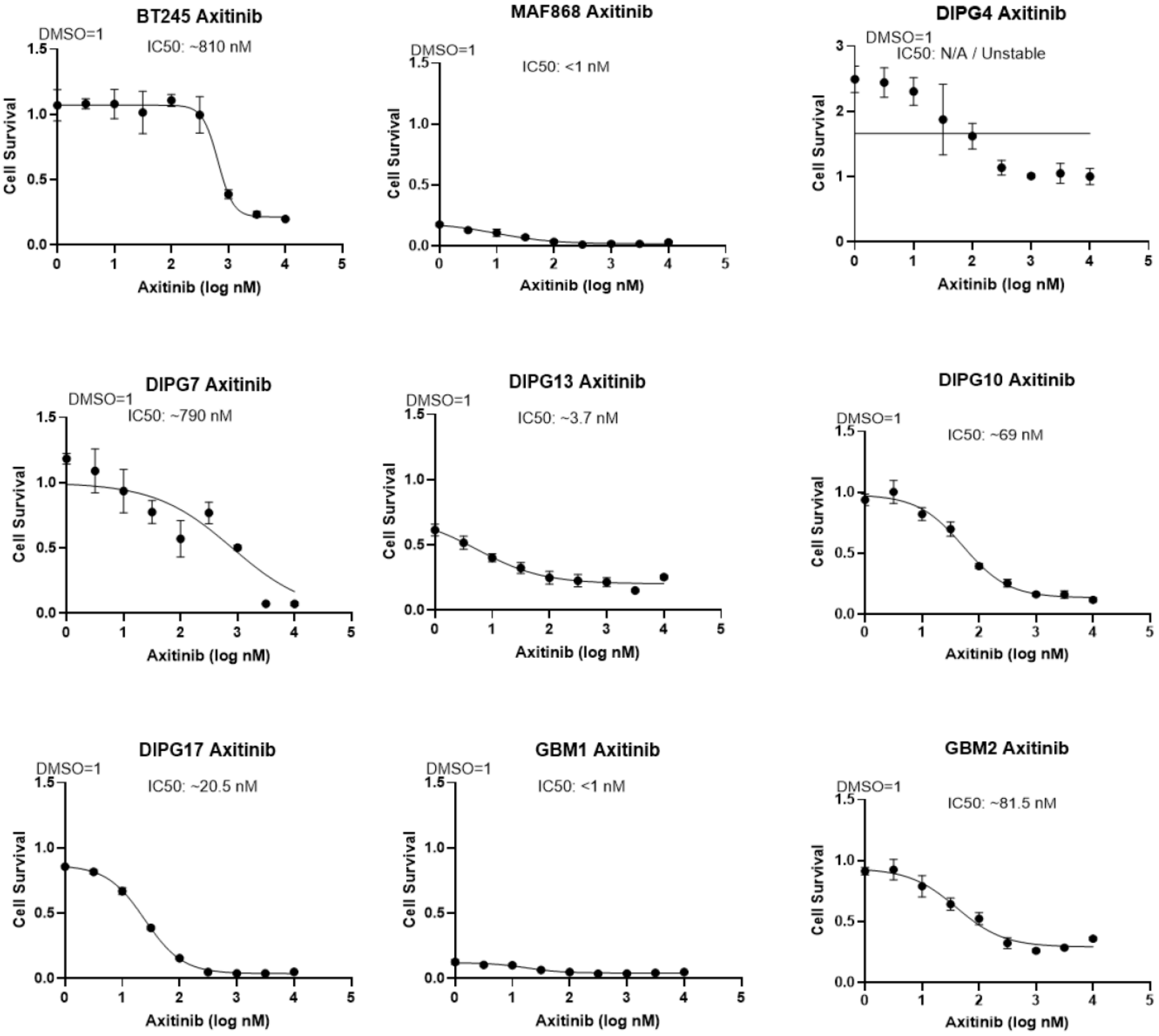

**Supplemental Figure 9:** *Dose-response curves for axitinib in pediatric glioma cell lines.*

Cell viability was measured by MTS assay following treatment with increasing concentrations of axitinib. Curves were fit using nonlinear regression (four-parameter logistic model) in GraphPad Prism. Error bars represent mean ± SD where replicate measurements were available. Estimated $IC_{50}$ values are shown within each panel.

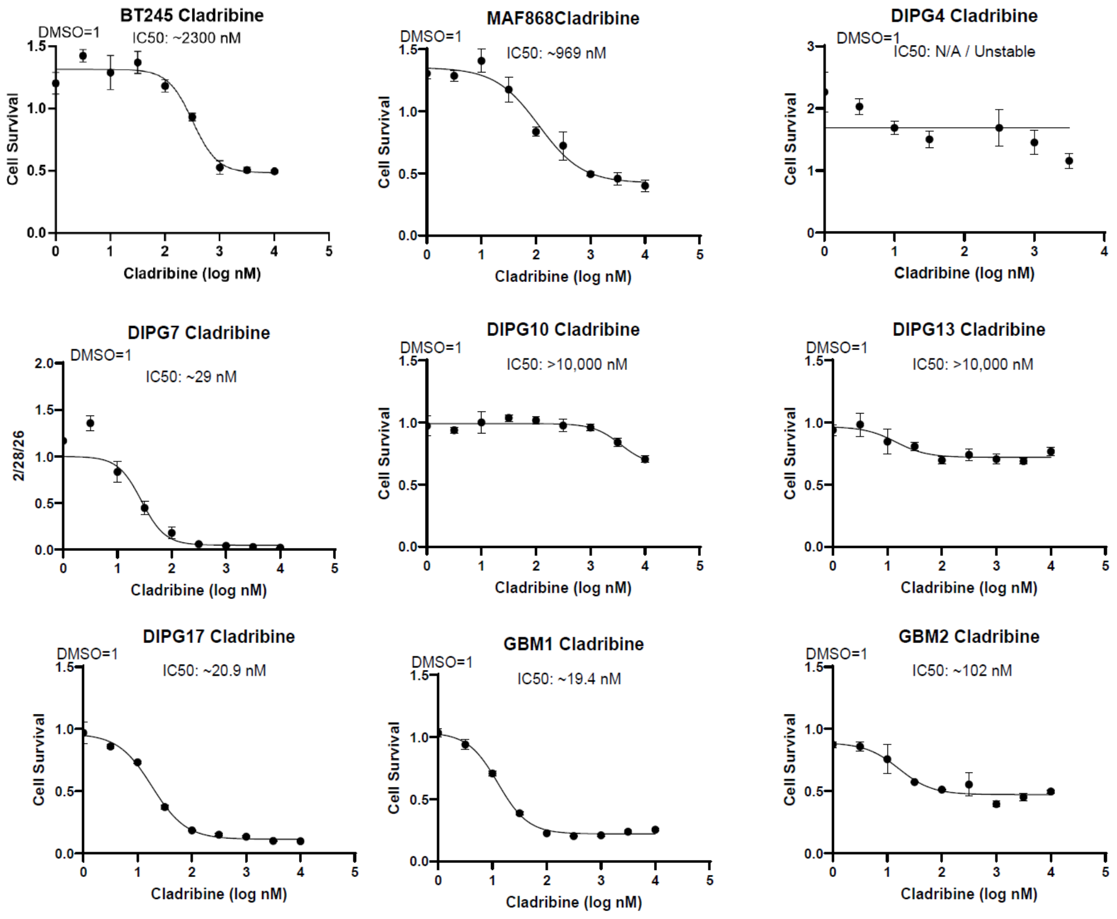

**Supplemental Figure 10:** *Dose-response curves for cladribine in pediatric glioma cell lines.*

Cell viability was measured by MTS assay following treatment with increasing concentrations of cladribine. Curves were fit using nonlinear regression (four-parameter logistic model) in GraphPad Prism. Error bars represent mean ± SD where replicate measurements were available. Estimated $IC_{50}$ values are shown within each panel.

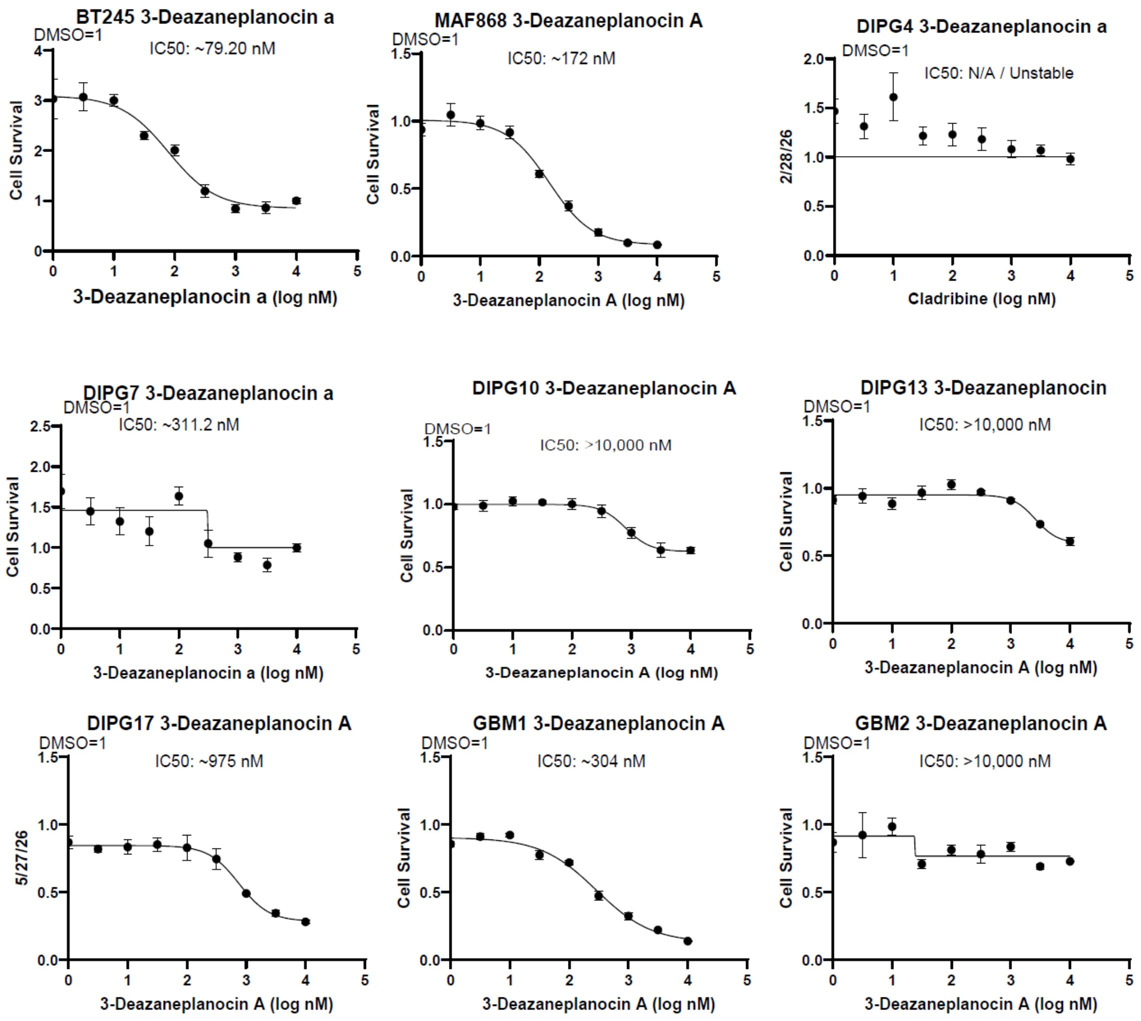

**Supplemental Figure 11:** *Dose-response curves for 3-Deazaneplanocin A in pediatric glioma cell lines.*

Cell viability was measured by MTS assay following treatment with increasing concentrations of 3-deazaneplanocin a. Curves were fit using nonlinear regression (four-parameter logistic model) in GraphPad Prism. Error bars represent mean ± SD where replicate measurements were available. Estimated $IC_{50}$ values are shown within each panel.
